# Detecting distortions of peripherally-presented letter stimuli under crowded conditions

**DOI:** 10.1101/048272

**Authors:** Thomas S. A. Wallis, Saskia Tobias, Matthias Bethge, Felix A. Wichmann

**Affiliations:** Neural Information Processing Group, Faculty of Science, Eberhard Karls Universität Tübingen, Werner Reichardt Center for Integrative Neuroscience, Eberhard Karls Universität Tübingen & the Bernstein Center for Computational Neuroscience, Tübingen; Neural Information Processing Group, Faculty of Science, Eberhard Karls Universität Tübingen; Werner Reichardt Center for Integrative Neuroscience, Eberhard Karls Universität Tübingen, Bernstein Center for Computational Neuroscience, Tübingen, Institute for Theoretical Physics, Eberhard Karls Universität Tübingen & the Max Planck Institute for Biological Cybernetics, Tübingen; Neural Information Processing Group, Faculty of Science, Eberhard Karls Universität Tübingen, Bernstein Center for Computational Neuroscience, Tübingen & the Max Planck Institute for Intelligent Systems, Empirical Inference Department, Tübingen

**Author notes:** Correspondence should be addressed to TSAW. TSAW was supported by an Alexander von Humboldt Postdoctoral Fellowship. Funded, in part, by the German Federal Ministry of Education and Research (BMBF) through the Bernstein Computational Neuroscience Program Tübingen (FKZ: 01GQ1002), the German Excellency Initiative through the Centre for Integrative Neuroscience Tübingen (EXC307), and the German Science Foundation (DFG; priority program 1527, BE 3848/2-1).

**Keywords:** 2D shape and form, spatial vision, reading, distortion, metamorphopsia

## Abstract

When visual features in the periphery are close together they become difficult to recognise: *something* is present but it is unclear what. This is called “crowding”. Here we investigated sensitivity to features in highly familiar shapes (letters) by applying spatial distortions. In Experiment 1, observers detected which of four peripherally-presented (8 deg of retinal eccentricity) target letters was distorted (spatial 4AFC). The letters were presented either isolated or surrounded by four undistorted flanking letters, and distorted with one of two types of distortion at a range of distortion frequencies and amplitudes. The bandpass noise distortion (“BPN”) technique causes spatial distortions in cartesian space, whereas radial frequency distortion (“RF”) causes shifts in polar coordinates. Detecting distortions in target letters was more difficult in the presence of flanking letters, consistent with the effect of crowding. The BPN distortion type showed evidence of tuning, with sensitivity to distortions peaking at approximately 6.5 c/deg for unflanked letters. The presence of flanking letters causes this peak to rise to approximately 8.5 c/deg. In contrast to the tuning observed for BPN distortions, RF distortion sensitivity increased as the radial frequency of distortion increased. In a series of follow-up experiments we found that sensitivity to distortions is reduced when flanking letters were also distorted, that this held when observers were required to report which target letter was undistorted, and that this held when flanker distortions were always detectable. The perception of geometric distortions in letter stimuli is impaired by visual crowding.

---

When a target object (such as a letter) is presented to the peripheral retina flanked by similar non-target objects (other letters), a human observer’s ability to discriminate or identify the target object is impaired relative to conditions where no flankers are present. This “crowding” phenomenon (Andriessen & Bouma, 1975; Bouma, 1970; Greenwood, Bex, & Dakin, 2009; Harrison & Bex, 2015; Herzog, Sayim, Chicherov, & Manassi, 2015; Levi, Klein, & Aitsebaomo, 1985; Parkes, Lund, Angelucci, Solomon, & Morgan, 2001; Strasburger, 2014; Toet & Levi, 1992) is characterised by a reduction in sensitivity to peripheral image structure. One way to physically change image structure is to apply spatial distortion, in which the position of local elements (pixels) are perturbed in some fashion (for example, by stretching or shifting). Characterising human sensitivity to spatial distortions is one way to investigate the perceptual encoding of local image structure. For example, showing that perception is invariant to a certain type of distortion (i.e. things look the same whether physically distorted or not) implies that the human visual system does not encode the distortion in question, either directly or indirectly. Arguably, measuring sensitivity to the distortion of highly familiar shapes such as letters (as we do in this paper) allows one to characterise human perception in a more complex task than (for example) grating orientation discrimination, but one that is more tractable from a modelling perspective than (for example) letter identification, which may require a full model of letter encoding. In addition, psychophysical investigation of spatial distortions is relevant to metamorphopsia—the perception of persistent spatial distortions in everyday life—which is commonly associated with retinal diseases that affect the macular (Wiecek, Dakin, & Bex, 2014).

Human sensitivity to spatial distortions has been investigated previously in images of faces (Dickinson, Almeida, Bell, & Badcock, 2010; Hole, George, Eaves, & Rasek, 2002; Rovamo, Mäkelä, Näsänen, & Whitaker, 1997; Spence, Storrs, & Arnold, 2014) and natural scenes (Bex, 2010; Kingdom, Field, & Olmos, 2007). To our knowledge, only one study has assessed the impact of spatial distortion for letter stimuli. Wiecek et al. (2014) had observers identify letters (26-alternative identification task) distorted with bandpass noise distortion (see below) while varying the spatial scale of distortion, the letter size and the viewing distance. Interestingly, they report an interaction between the spatial scale of distortion (CPL; cycles per letter) and viewing distance (changing letter size), such that for small letters (subtending 0.33 degrees of visual angle) performance was worst for coarse-scaled distortions (2.4 CPL), whereas for large letters (5.4 deg) the most detrimental distortion shifted to a finer scale (4 CPL). This result has important implications for patients with metamorphopsia: a stable retinal distortion may affect letter recognition for some letter sizes but not others, influencing acuity assessments using letter charts (a primary outcome measure for clinical vision assessment; Wiecek et al., 2014).

Here we investigate sensitvity to spatial distortions in letters, under crowded (flanked) and uncrowded (unflanked) conditions. Note that our goal here is distinct from that of Wiecek et al. (2014), who measured the impact of distortions on letter identification. We do not measure letter identification here, but instead use letters as a class of relatively simple, artifical, but highly familiar stimuli to investigate sensitivity to the presence of distortion *per se.* We quantify the detectability of two different types of spatial distortion commonly used in the literature (see also Stojanoski & Cusack, 2014, for another distortion not employed here). In bandpass noise distortions (hereafter referred to as *BPN* distortion; Bex, 2010), pixels are warped according to bandpass filtered noise; this ensures that the distortion occurs on a defined and limited spatial scale. In radial frequency distortions (hereafter referred to as *RF* distortion; Dickinson et al., 2010; Wilkinson, Wilson, & Habak, 1998), the image is warped by modulating the radius (defined from the image centre) according to a sinusoidal function of some frequency defined in polar coordinates. For our purposes they serve to produce two different graded changes in letter images. A successful model of form discrimination in humans would explain sensitivity to both types of distortion and any dependence on surrounding letters (potentially, different mechanisms may be required to explain sensitivity to each distortion type).

## Experiment 1

### Methods

Stimuli, data and code associated with this paper are available to download from http://dx.doi.org/10.5281/zenodo.159360. This document was prepared using the `knitr` package (Xie, 2013, 2015) in the R statistical environment (Arnold, 2016; Auguie, 2016; R Core Development Team, 2016; Wickham, 2009, 2011; Wickham & Francois, 2016) to increase its reproducibility.

#### Observers

Five observers with normal or corrected-to-normal vision participated in this experiment: two of the authors, one lab member and two paid observers (10 Euro per hour) who were unaware of the purpose of the study. All of the observers had prior experience with psychophysical experiments and were between 20 an 31 years of age. All experiments conformed to Standard 8 of the American Psychological Association’s Ethical Principles of Psychologists and Code of Conduct (2010).

#### Apparatus

Stimuli were displayed on a VIEWPixx LCD (VPIXX Technologies; spatial resolution 1920×1200 pixels, temporal resolution 120 Hz). Outside the stimulus image the monitor was set to mean grey. Observers viewed the display from 60 cm (maintained via a chinrest) in a darkened chamber. At this distance, pixels subtended approximately 0.024 degrees on average (41.5 pixels per degree of visual angle). The monitor was carefully linearised (maximum luminance 212 cd/m^2^) using a Gamma Scientific S470 Optometer. Stimulus presentation and data collection was controlled via a desktop computer (12 core i7 CPU, AMD HD7970 graphics card) running Kubuntu Linux (14.04 LTS), using the Psychtoolbox Library (Brainard, 1997; Kleiner, Brainard, & Pelli, 2007; Pelli, 1997, version 3.0.11) and our internal iShow library (http://dx.doi.org/10.5281/zenodo.34217) under MAT-LAB (The Mathworks, Inc., R2013B). Responses were collected using a RESPONSEPixx button box.

#### Stimuli

The letters stimuli were a subset of the Sloan alphabet (Sloan, 1959), used commonly on acuity charts to measure visual acuity in the clinic. Target letters were always the letters D, H, K and N; flanker letters were always C, O, R, and Z. Letter images were 64 × 64 pixels. To prevent border artifacts in distortion, each image was padded with white pixels of length 14 at each side, creating 92 × 92 pixel images. These padded letter images were distorted according to distortion maps generated from the BPN or RF algorithms (see below) in a Python (v2.7.6) environment, using Scipy’s `griddata` function with linear 2D interpolation to remap pixels from the original to the distorted image. That is, the distortion map specifies where to move the pixels from the original image; pixel values in intermediate spaces are linearly interpolated from surrounding pixels to produce smooth distortions.

#### Bandpass Noise (BPN) distortion

Bex (2010, see also (Rovamo et al., 1997; Wiecek et al., 2014)) describes a method for generating spatial distortions that are localised to a particular spatial passband (see Figure 1A–D). Two random 92 × 92 samples of zero-mean white noise were filtered by a log exponential filter (see Equation 1 in Bex, 2010):

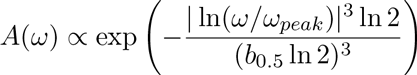

where *ω*_*peak*_ specifies the peak frequency, *ω* is the spatial frequency and b_0.5_ is the half bandwidth of the filter in octaves. Noise was filtered at one of six peak frequencies (2, 4, 6, 8, 16, 32 cycles per image; corresponding to 1.3, 2.6, 4, 5.3, 10.6 and 21.3 c/deg under our viewing conditions) with a bandwidth of one octave. The filtered noise was windowed by multiplying with a circular cosine of value one, falling to zero at the border over the space of 14 pixels, ensuring that letters did not distort beyond the borders of the padded image region. The amplitude of the filtered noise was then rescaled to have max / min values at 0.25, 0.5, 1, 1.5, 2, 2.5, 3, or 5 pixels; this controlled the strength of the distortion. For presentation of the results (thresholds, below), these amplitude units were transformed from pixels to degrees. One filtered noise sample controlled the horizontal pixel displacement, the other controlled vertical displacement (together giving the distortion map for the `griddata` algorithm).

**Figure 1.**
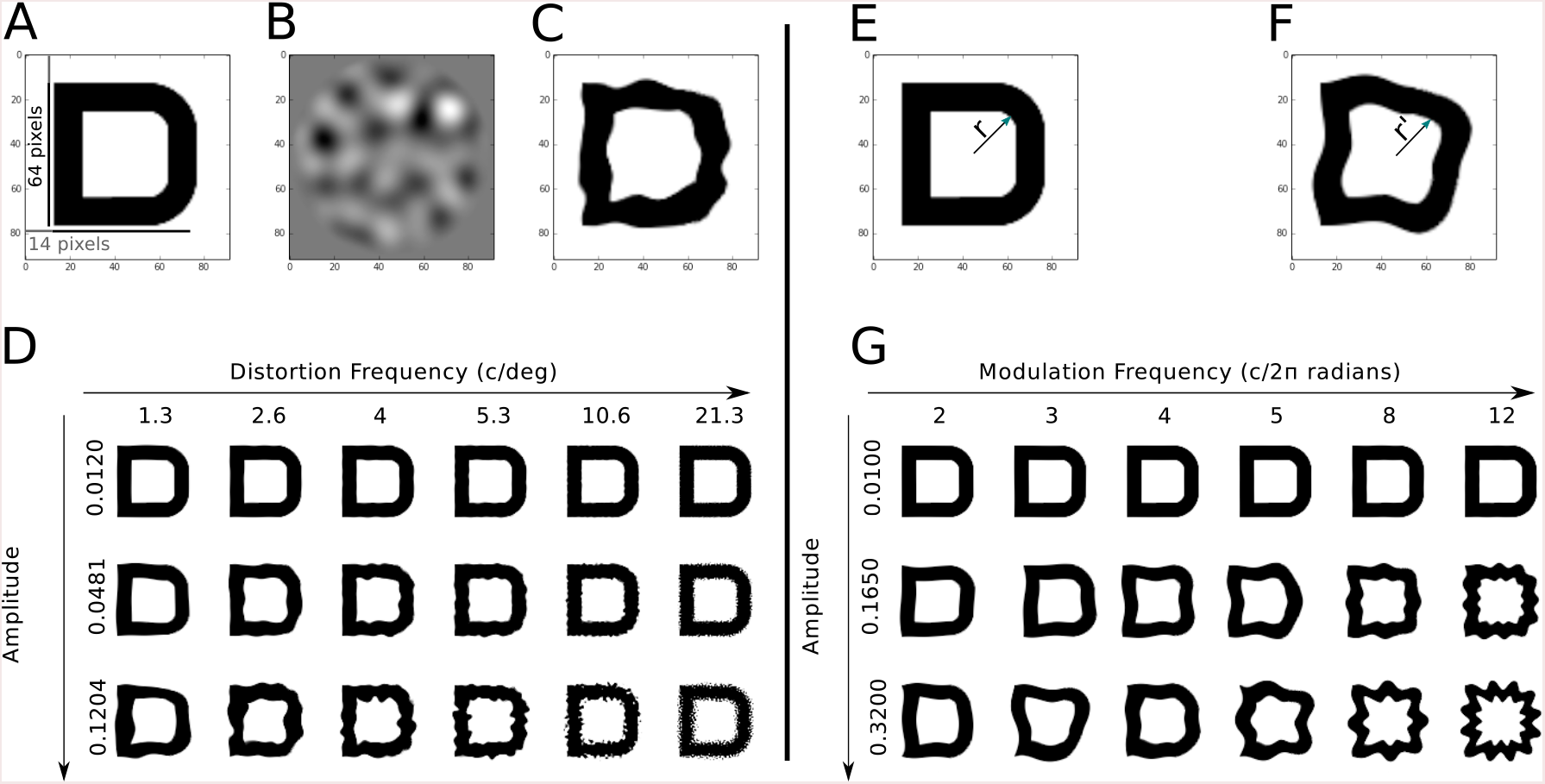
Distortion methods for Bandpass Noise (BPN; A–D) and Radial Frequency (RF; E–G). **A:** A Sloan letter (D) with 14 pixels of white padding. **B:** A sample of bandpass filtered noise, windowed in a circular cosine. Two such noise samples determine the BPN distortion map. **C:** The letter distorted by the BPN technique. **D:** The effects of varying the frequency (columns) and amplitude (rows) of the BPN distortion. **E:** An original letter image, showing the original radius *r* from the centre to an arbitrary pixel. **F:** RF distortion modulates the radius of every pixel according to a sinusoid, producing a new radius *r* ′. **G:** The effects of varying the frequency (columns) and amplitude (rows) of the RF distortion. More examples of distortions applied to letters are provided in the Supplementary Material.

#### Radial Frequency (RF) distortion

Here, the distortion map was created by modulating the distance of each pixel from the centre of the padded image according to a sinusoid defined in polar coordinates (see Equation 3 in Wilkinson et al., 1998, and 1E–G):

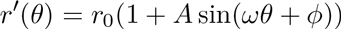

where *r* ′ is the distorted radius from the centre, *r*_*o*_ the undistorted (mean) radius, *A* is the amplitude of distortion (the proportion of the unmodulated distance from the centre), *θ* is the polar angle and *ω* is the radial frequency of distortion (here 2, 3, 4, 5, 8 or 12 cycles in 2*π* radians). The angular phase of the modulation (*ϕ*) on each trial was drawn from a random uniform distribution spanning [0, 2*π*]. The amplitude of the distortion was set to one of 0.0075, 0.01, 0.0617, 0.1133, 0.1650, 0.2167, 0.2683 or 0.3200. The distortion map was windowed in a circular cosine as above, then the cosine and sine values were passed to `griddata` as the horizontal and vertical offsets.

To facilitate future modelling of our experiment, we pregenerated all images presented to observers (see below) and saved them to disk. In total we generated 1920 images: two distortion types (BPN, RF) × two conditions (flanked, unflanked; see below) × eight amplitudes × six frequencies, each repeated 10 times. BPN distortions are generated from new random noise images and RF distortions with random phases, meaning that these 10 repetitions were unique images. Target positions, letter identities and distortions were randomised on each repeat. In addition, we generated the same 1920 images *without* applying distortion to one of the target letters and saved them to disk. An image-based model of pattern recognition could be evaluated on the same stimuli as we have shown to our observers, using an undistorted “full-reference” image as a baseline (all images are provided online at http://dx.doi.org/10.5281/zenodo.159360).

#### Procedure

On each *unflanked* trial, observers saw the four target letters and indicated the location (relative to fixation) of the distorted letter. The letters subtended approximately 1.5 × 1.5 dva and were located above, below, right and left of fixation (see Figure 2A); letter identity at each location was randomly shuffled on each trial. The target letters were centred at a retinal eccentricity of 320 pixels (7.7 dva), and observers were instructed to maintain fixation on the central fixation cross (best for steady fixation from Thaler, Schütz, Goodale, & Gegenfurtner, 2013). The entire letter array was presented on a square background of maximum luminance (side length 1024 pixels or 24.3 dva); the remainder of the monitor area was set to mean grey. Letter strokes were set to minimum luminance (i.e. the letters were approximately 100% Michelson contrast). The letter array was presented for 150 ms (abrupt onset and offset), after which the screen was replaced with a fixation cross on the same square bright background. The observer had up to 2000 ms to respond (a response triggered the next trial with ITI 100 ms), and received auditory feedback as to whether their response was correct.

**Figure 2.**
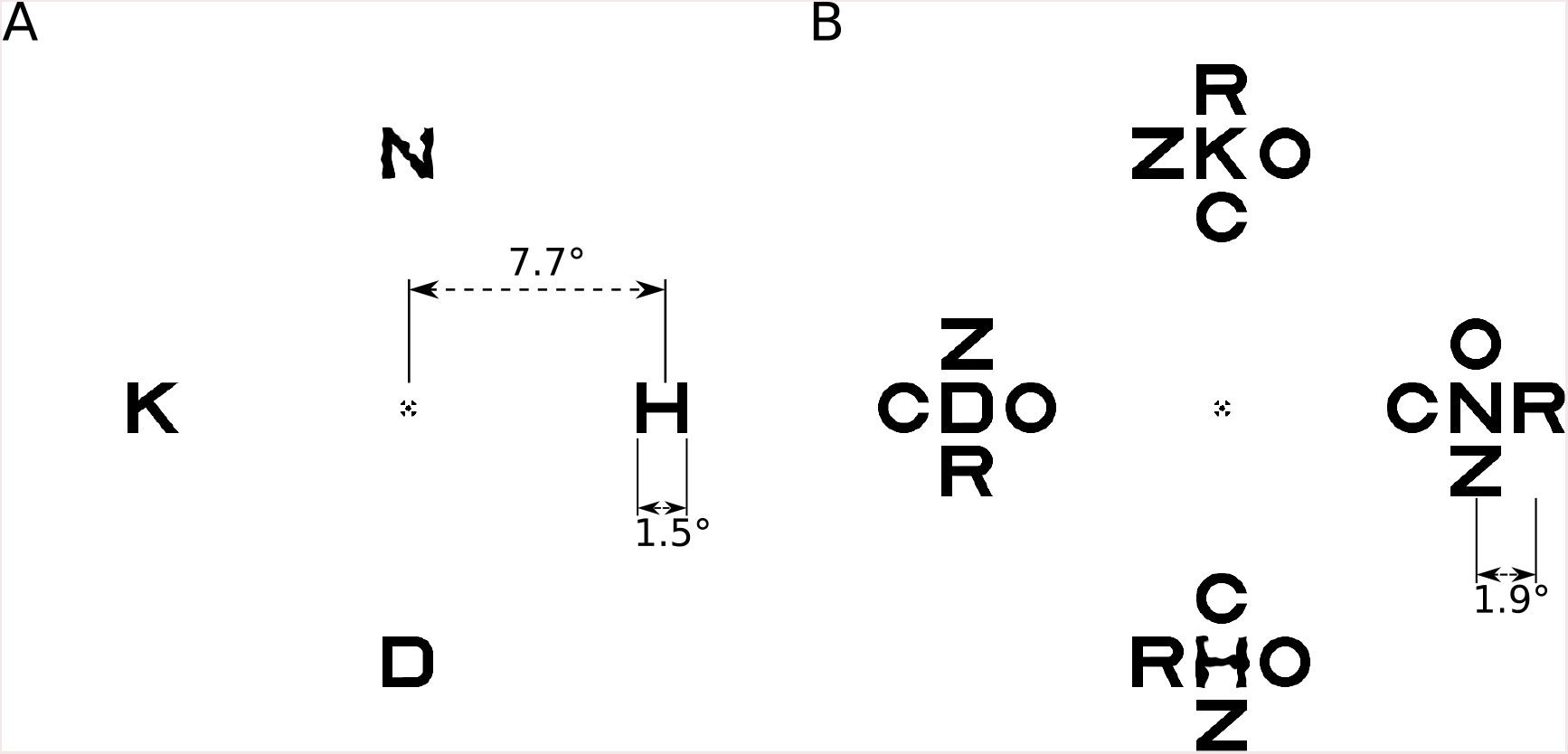
Example stimulus arrays showing BPN distortions. **A:** An unflanked trial example. In this example the correct response is “above”. **B:** A flanked trial example. The correct response is “below”.

On *flanked* trials (Figure 2B), four undistorted flanking letters the same size as the target were presented above, below, left and right of each target letter (centre-to-centre separation 1.9°, corresponding to approximately 0.25 of the eccentricity, well within the spacing of “Bouma’s law”; Bouma (1970)). The arrangement of the four flanking letters was randomly determined on each trial.

Different distortion frequencies (six levels) and amplitudes (seven levels^1^) were randomly interleaved within a block of trials, whereas the distortion type (BPN or RF) and letter condition (unflanked or flanked) were presented in separate blocks. Each pairing of frequency and amplitude was repeated 10 times (corresponding to the unique images generated above), creating 420 trials per block. Breaks were enforced after every 70 trials. Blocks of trials were arranged into four-block sessions, in which observers completed one block of each pairing of distortion type and letter condition. Observers always started the session with an unflanked letter condition in order to familiarise them with the task ^2^. Each session took approximately two hours. All observers participated in at least four sessions. Before the first block of the experiment observers completed 70 practice trials to familiarise themselves with the task. In total we collected 20,160 trials on each of the unflanked and flanked conditions.

#### Data analysis

Data from each experimental condition were fit with a cumulative Gaussian psychometric function using the *psignifit 4* toolbox for Matlab (Schütt, Harmeling, Macke, & Wichmann, 2016), with the lower asymptote fixed to chance performance (0.25). The posterior mode of the threshold parameter (midpoint of the unscaled cumulative function) and 95% credible intervals were calculated using the default (weak) prior settings from the toolbox. The 95% credible intervals mean that the parameter value has a 95% probability of lying in the interval range, given the data and the prior. Psychometric function widths (slopes) either did not vary appreciably over experimental conditions (Experiment 1) or, when they did (Experiment 2), patterns of variation showed effects consistent with the threshold estimates. This paper therefore presents only threshold data for brevity.

### Results

Thresholds for detecting the distorted target letter are shown in Figure 3. For both distortion types, observers were less sensitive to letter distortion (thresholds were higher) when the target letters were surrounded by four flanking letters (light triangles) compared to when targets were isolated (dark circles). This pattern is an example of crowding. Furthermore, we observe that the two distortion types (BPN and RF) show different dependencies on their respective frequency parameters (which are not themselves comparable). RF distortions become easier to detect the higher their frequency (c / 2*π* radians). BPN distortions show evidence of tuning, such that thresholds are lowest for frequencies in the range of 4–10 c/deg and rise for both lower and higher frequencies (note the log-log scaling in Figure 3). To quantify these effects, we fit curves to the thresholds as a function of the log distortion frequency (BPN: four-parameter Gaussian fit by minimising the sum of squared errors with the BFGS method of R’s `optim` function^3^; RF: linear model fit with R’s `lm` function; see lines in Figure 3 for model fits).

**Figure 3.**
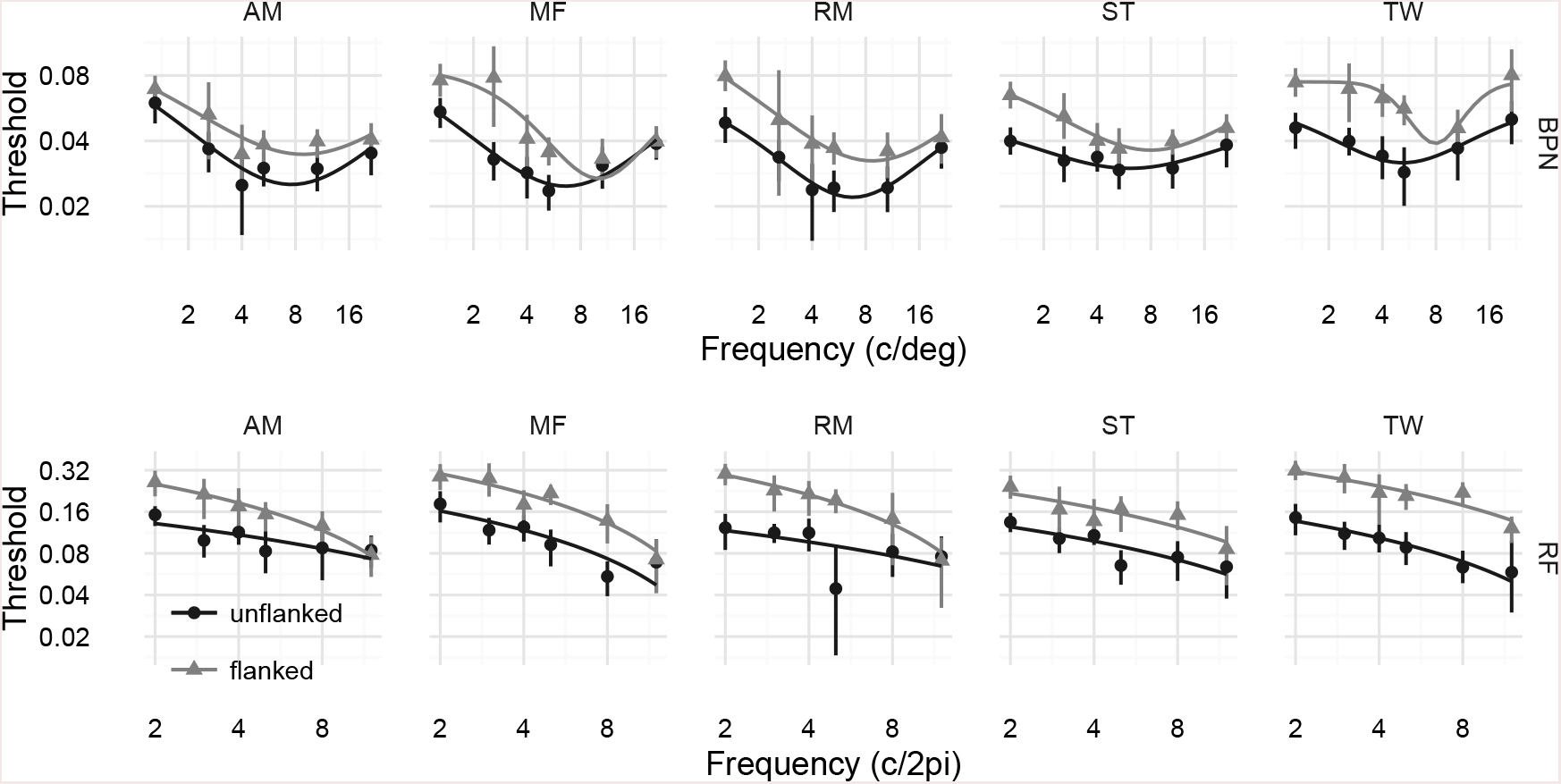
Results of Experiment 1. Top panels show threshold amplitude for detecting letters distorted with BPN distortions, as a function of distortion frequency (c/deg) for five observers. Note both the x-and y-axes are logarithmic. Points show the posterior MAP estimate for the psychometric function threshold; error bars show 95% credible intervals. Thresholds are higher (observers are less sensitive to distortions) when flanking letters are present (light triangles) compared to unflanked conditions (dark circles). Additionally, thresholds appear to show tuning, being lowest at approximately 6–8 c/deg. Lines show fits of a Gaussian function to the log frequencies and linear thresholds (see text for details). Bottom row of panels show RF distortions. Flanking letters again impair performance. Unlike in the BPN distortions, for RF distortions performance simply worsens for higher distortion frequencies. Lines show fits of a linear model to the log frequencies and linear thresholds. The reader can appreciate these results for themselves by examining how distortion visibility changes as a function of frequency in Figure 1D and G.

To quantify the overall decrease in performance caused by the presence of flanking letters, we examined how the area under these curves (estimated numerically) changed from unflanked to flanked conditions^4^. Larger areas mean higher thresholds (i.e. lower sensitivity). We quantify these differences using paired t-tests of both frequentist and Bayesian (Morey & Rouder, 2015; Rouder, Morey, Speckman, & Province, 2012) flavours. For the BPN distortion type, flanking letters raised the mean area under the Gaussian threshold curve from 0.09 (SD = 0.01) to 0.14 (SD = 0.02); t(4) = 6.26, p = 0.0033, BF = 15.7. For the RF distortion type, flanking letters raised the mean area under the linear fit from 0.17 (SD = 0.01) to 0.33 (SD = 0.05); t(4) = 7.17, p = 0.002, BF = 22.6. Thus both crowding effects we observe appear reasonably robust.

Next we consider the peak distortion frequency at which thresholds were lowest for the BPN distortions (there is no peak in our data for the RF distortions). There was a reasonable effect of flanking, such that when flanking letters were present, distortion sensitivity peaked at higher frequencies (M = 8.73 c/deg, SD = 0.88) than when target letters were unflanked (M = 6.44, SD = 0.88; a difference in peaks of 0.44 octaves; t(4) = 5.9, p = 0.0041, BF = 13.4). While the effect is therefore large compared to the relevant error variance, note that it ignores the precision with which the peak frequency is determined by the data, and so should be interpreted with a degree of caution.

## Experiment 2

Our first experiment showed that sensitivity to both BPN and RF distortions was reduced in the presence of undistorted flanking letters. Interestingly, our observers reported experiencing “pop-out” in the flanked condition, such that the distorted letter appeared relatively more salient than the three undistorted targets by virtue of its contrast with neighbouring undistorted flankers. That is, the distorted letter strokes appeared subjectively more noticable when next to undistorted strokes. While the data quantitatively argue against such a pop-out effect (since flanking letters impaired performance), we nevertheless decided to conduct a series of follow-up experiments to determine whether there was any dependence of the thresholds on the kind of flankers employed. Flankers more similar to the target are known to cause stronger crowding (e.g. Bernard & Chung, 2011; Kooi, Toet, Tripathy, & Levi, 1994); it is therefore plausible that distorted flankers would produce even greater performance impairment.

We test this hypothesis in three related sub-experiments. Because we will directly compare the data from each experiment, we present the similarities and differences in the experimental procedures first, followed by all data collectively. Three of the observers from Experiment 1 (two authors plus one lab member) participated in these experiments; all other experimental procedures were as in Experiment 1 except as noted below. As in Experiment 1, all test images were pregenerated and saved along with undistorted reference images to facilitate future modelling work.

### Methods

#### Experiment 2a: varying the number of distorted flankers

This experiment was identical to Experiment 1, with the primary exception that in some trials either two or four of the flanker letters in every letter array (above, left, below and right) were also distorted (see Figure 4A–C). That is, observers reported the location of the distorted target letter, sometimes in the presence of distorted flankers. If distorted targets pop out from undistorted flankers *and* undistorted targets pop out from distorted flankers *(symmetrical popout*), we might expect that settings in which two of four flankers are distorted would be hardest. In the case of no undistorted flankers (i.e. the same as the flanked condition in Experiment 1), the distorted target pops out from the flankers. In the case of four distorted flankers, the *un*distorted targets pop out in three of the four possible locations, alerting the observer to the correct response by elimination. Finally, when two flanking letters are distorted, any differential pop-out signal is minimised because the nontarget letter arrays contain two distorted letters whereas the letter array corresponding to the correct response contains three distorted letters. This account would therefore predict that thresholds in the two distorted flanker letter condition should be higher than those for zero or four distorted flankers.

**Figure 4.**
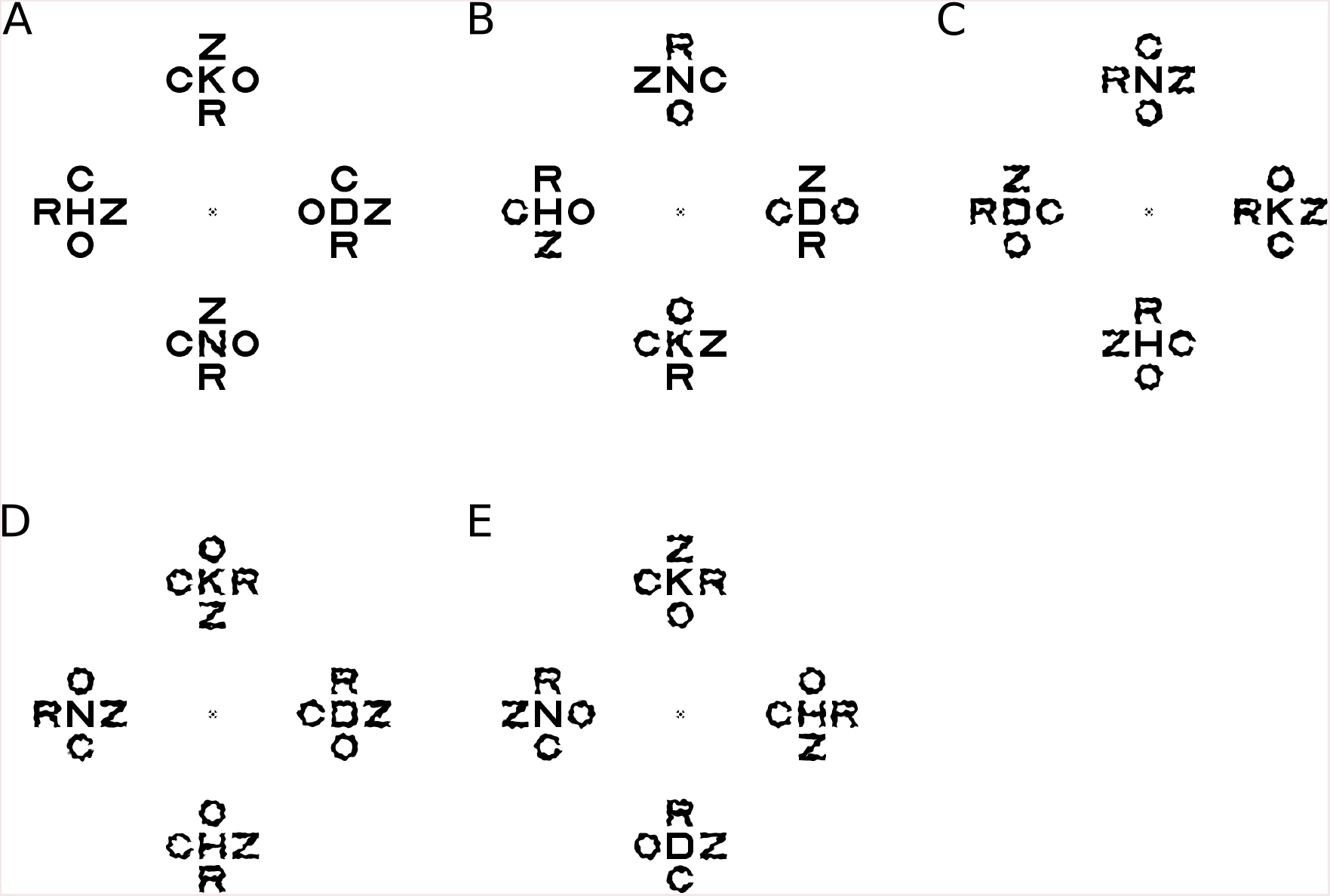
Example stimulus displays from Experiment 2 (all examples show the BPN distortion type at high distortion amplitudes). In Experiment 2a, observers detected the distorted middle letter when surrounded by zero **(A),** two **(B)** or four **(C)** distorted flankers. **D:** In Experiment 2b, observers indicated the *un*distorted middle letter surrounded by four distorted flankers. **E:** In Experiment 2c, flankers were always distorted at a highly-detectable distortion level. The correct response in panels A–E are down, left, down, left and right.

In this experiment we selected one distortion frequency for each distortion type: 2.6 c/deg for the BPN and 4 c/2*π* for the RF distortions. Because our pilot testing indicated these tasks were more difficult than those in Experiment 1, we generated distortions at higher amplitudes than those in the first experiment: 0.024, 0.048, 0.072, 0.096, 0.120, 0.144, and 0.168 for BPN and 0.05, 0.125, 0.2, 0.275, 0.25, 0.425 and 0.5 for RF. Flanking letters were distorted with the same frequency and amplitude distortion as the target letter on every trial.

Trials of different distortion types (BPN, RF) and flanker conditions (zero, two or four distorted flankers) were presented in separate blocks in which each of the seven amplitudes were randomly interleaved. Ten unique images were created for each amplitude, each repeated three times to give 30 trials per amplitude (210 per block). Blocks of trials were arranged into six-block sessions, consisting of each distortion type and flanker condition in a random order for each observer. All observers participated two sessions, creating a total of 7560 trials.

#### Experiment 2b: detect the undistorted letter in the presence of distorted flankers

In Experiment 1, observers detected which of four letters was distorted when surrounded by four undistorted flanking letters. In Experiment 2b we examine the inverse task: to detect which middle letter is *un*distorted in the presence of four distorted flankers (Figure 4D). If distortion detection is symmetric, performance in this condition should be as good as in the zero distorted flanker condition of Experiment 2a. That is, distorted letters should pop out from undistorted flankers just as undistorted letters pop out from distorted flankers. The procedure was otherwise identical to Experiment 2a, with the exception that observers did two blocks (BPN and RF distortion types) of 210 trials (totalling 1260 trials).

#### Experiment 2c: flanker distortion at fixed high amplitude

In Experiments 2a and 2b, flanker distortions had the same amplitude as the target letter distortion. Therefore, for low target distortion amplitudes the flanker distortions were also subthreshold. Popout, if it exists, may require detectable levels of distortion in the flanking elements. To test this question we repeated the four distorted flanker condition from Experiment 2a, with the exception that the flankers were distorted at a fixed amplitude that rendered distortions easily detectable (0.144 c/deg for BPN, 0.425 c/2*π* for RF; see Figure 4E). If popout requires suprathreshold distortions in flanking letters then sensitivity in this condition should be higher than the four distorted flanker condition from Experiment 2a (i.e. more similar to the zero distorted flanker condition for Experiment 2a). Observers performed at least two blocks, one for each distortion type (2520 trials total).

### Results

Threshold levels of distortion are shown in Figure 5. The results for the BPN and RF distortions show qualitatively similar effects of the experimental conditions. First, thresholds increase as more flanking letters are distorted: detecting distortions in arrays with two or four distorted flankers is more difficult than when no flankers are distorted (Experiment 2a; Figure 5 circles). There is therefore no support for the prediction that thresholds would be higher in the two distorted flanker condition which, had it occurred, would be consistent with targets popping out from (un)distorted flankers in the zero and four distorted flanker conditions.

**Figure 5.**
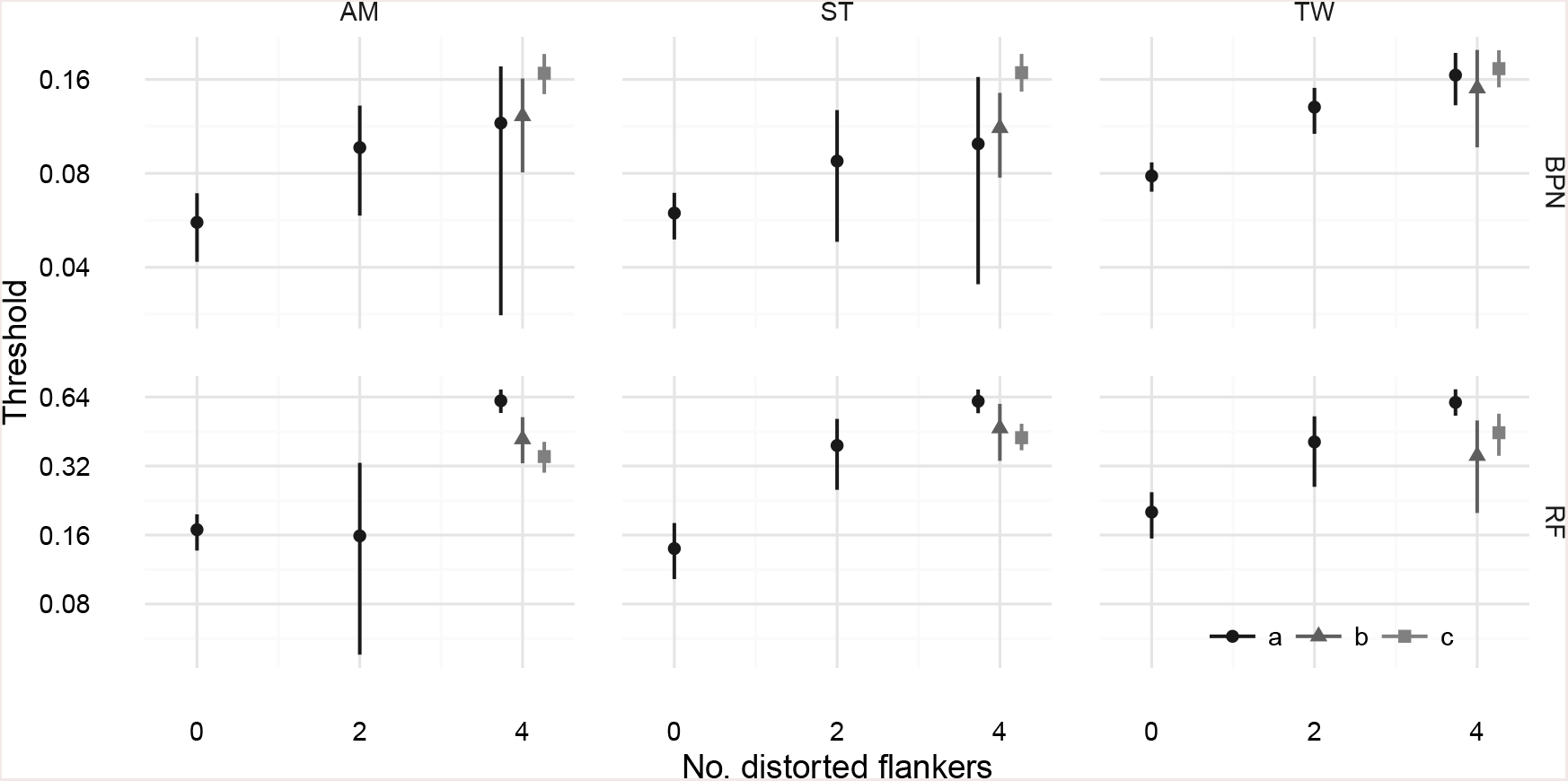
Results of Experiment 2. Top panels show threshold amplitude for detecting the target letter as a function of the number of distorted flankers, for three observers in the BPN distortion condition (Experiment 2a). Note the logarithmic y-axis. Points show the posterior MAP estimate for the psychometric function threshold; error bars show 95% credible intervals. Points for four distorted flankers have been shifted in the x direction to aid visibility. Bottom panels show the same as the top for RF distortions.

The results of Experiment 2b (Figure 5, triangles) also provided no support for symmetrical popout. There was no evidence that detecting an undistorted target letter amongst four distorted flankers was as difficult as the zero distorted flanker condition of Experiment 2a; instead, thresholds for detecting the undistorted target letter were more similar to those for detecting a distorted target letter amongst four distorted flankers.

Finally, thresholds in Experiment 2c (Figure 5, squares) show that detecting a distorted letter amongst four distorted flankers requires substantially more distortion amplitude than those with no distorted flankers (Experiment 2a with no distorted flankers), despite the flanker distortions always being easily detectable. This result confirms the absence of symmetrical popout found in Experiments 2a and 2b: it is not the case that the three undistorted targets pop out from their distorted surrounds (which if it occurred would allow the observer to choose the correct response by selecting the array with no popout).

It is additionally interesting to consider the pattern of results for Experiment 2c relative to the other four letter distorted flanker conditions. Here we see opposite patterns of results for the BPN and RF distortions. For BPN distortions, Experiment 2c produces the highest thresholds compared to the other experiments, suggesting that highly visible flanker distortions produce even stronger masking. Conversely, for the RF distortions Experiment 2c thresholds are lowest of the other four-distorted-flanker data in two of three observers. This could reflect some facilitation for this distortion type, but given the inconsistency between observers we would want to collect more data before drawing strong conclusions.

## Discussion

We have measured human sensitivity to geometric distortions of letter stimuli presented to the peripheral retina. For two types of distortion, Experiment 1 showed that distortion sensitivity is reduced when target letters are surrounded by task-irrelevant flankers. This result is therefore an example of crowding (Bouma, 1970). In the follow-up studies of Experiment 2 we found that this impairment became more severe^5^ when flanking letters were themselves distorted – i.e. we do not find evidence of distortion “pop-out”. That distortion sensitivity can be crowded is perhaps unsurprising; nevertheless, we find it worthwhile to demonstrate the impairment and measure its strength. The second result is more curious, because a consideration of the stimulus dimensions that may underlie distortion detection suggests we should have found the opposite result.

### Relevance to crowding

Crowding has previously been shown to exist for both letter identification (Bouma, 1970; Chung, Legge, & Tjan, 2002; Estes, 1982; Pelli, Palomares, & Majaj, 2004) and orientation discrimination (Andriessen & Bouma, 1975; Harrison & Bex, 2015; Parkes et al., 2001; Pelli et al., 2004; Wilkinson, Wilson, & Ellemberg, 1997). Our experiments could be considered to probe an intermediate level of representation: geometric distortions can change the contours of these simple but highly familiar shapes.

It is therefore relevant to ask what more primitive dimensions might underlie the effects we report. Detecting deviations from expected shape potentially involves local orientation processing, position, curvature, contour alignment and spatial frequency changes. What does the crowding literature tell us about these potential cues? As mentioned above, there is strong evidence from a number of studies that local orientation processing is impaired by crowding. Sensitivity to local position (Dakin, Cass, Greenwood, & Bex, 2010; Greenwood et al., 2009; Greenwood, Bex, & Dakin, 2012), spatial frequency (Wilkinson et al., 1997), curvature (Kramer & Fahle, 1996), and contour alignment (Chakravarthi & Pelli, 2011; Dakin & Baruch, 2009; May & Hess, 2007; Robol, Casco, & Dakin, 2012) is also impaired by flanking elements. Some or all of these potential cues could therefore be related to the effects we observe.

The results from our second experiment show that distorted targets do not pop out from undistorted flankers (and vice versa). This is interesting in light of the extensively-documented effects of target-flanker similarity in crowding (Bernard & Chung, 2011; Chakravarthi & Pelli, 2011; Chung, Levi, & Legge, 2001; Estes, 1982; Glen & Dakin, 2013; Herzog et al., 2015; Kooi et al., 1994; Livne & Sagi, 2007, 2010; Manassi, Sayim, & Herzog, 2013; Saarela, Sayim, Westheimer, & Herzog, 2009; Sayim & Cavanagh, 2013; Wilkinson et al., 1997). If we define “similarity” at the level of “distortedness”, then in Experiment 1 the distorted target becomes less similar to the undistorted flankers as distortion amplitude increases. The degree of target-flanker similarity in the non-target letter arrays is constant, and determined only by the confusability of the undistorted letters in those arrays. The same holds true for Experiment 2a in the zero distorted flankers condition. When the four flankers are also distorted in Experiment 2a, the similarity between target and flankers in the target array is held constant (as the target becomes distorted with increasing amplitude, so do the flankers), whereas in the non-target letter arrays the central (undistorted) letters and the distorted flankers become less similar. If observers were able to use this decreasing similarity to rule out the non-target arrays, we would expect them to be sensitive to the target location. Instead their thresholds are much higher relative to the zero distorted flankers case Experiment. 2c provides the opposite case to Experiment 1: because flankers were distorted with a strong amplitude distortion, then as distortion amplitude in the target letter increases, it becomes more similar to the flankers. Therefore, target-flanker similarity effects defined at the level of “distortedness” do not appear to be generally consistent with the patter of results we observe.

A more parsimonious account consistent with the results of Experiment 2 is that performance decays as the “complexity” of the stimulus array increases (under the assumption that flanker distortion increases complexity) ^6^. When all four flanking letters were distorted (Figure 5), thresholds for target detection were higher than other conditions whether the observers were trying to discriminate a distorted middle letter from undistorted ones (Experiment 2a), the undistorted middle letter from distorted middle letters (Experiment 2b) or the distorted middle letter in the presence of strong flanker distortions (Experiment 2c). Flanker distortion increases complexity, making the task more difficult. Letter complexity effects have indeed been demonstrated to play a distinct role from target-flanker similarity in crowded letter identification (Bernard & Chung, 2011), an effect attributed to the number of features to be detected within a character (see also Pelli, Burns, Farell, & Moore-Page, 2006; Suchow & Pelli, 2012). It seems plausible then that in our Experiment 2, it is difficult to detect distorted letters in the presence of distorted flankers because of feature crowding.

The model of letter complexity presented by Bernard and Chung (2011) requires a letter skeleton to be known (their paper compared different fonts). We require an image-based metric. We made a coarse attempt to quantify the complexity account above by investigating whether two metrics of visual clutter (Rosenholtz, Li, & Nakano, 2007) could qualitatively mimic the effects—on the assumption that a complex display is a cluttered display. Feature congestion is a multiscale measure of the covariance of the luminance contrast, orientation and colour in a given input image. *Subband* entropy is determined by the bitdepth required for wavelet image encoding, expressed as Shannon entropy in bits. These metrics have previously been associated with performance in tasks such as visual search (Asher, Tolhurst, Troscianko, & Gilchrist, 2013; Henderson, Chanceaux, & Smith, 2009; Rosenholtz et al., 2007). While both metrics showed a robust increase in clutter from unflanked to flanked displays, there was only weak evidence that they were able to capture the other effects in our data (see Supplementary Material). One would need to find a more appropriate measure of complexity—perhaps something similar to these clutter metrics—to capture the full range of the data we report.

Two dominant classes of crowding models are “averaging” models, in which crowding occurs because task-relevant features from the target and flankers are averaged together, and “substitution” models in which properties of the flankers are sometimes mistakenly reported as properties of the target. The present study was not designed to discriminate between these accounts of crowding, and it is somewhat unclear what predictions models of either class would make for our results (can the appearance of distortion be substituted?). Interestingly, recent work shows that because both averaging-and substitution-like errors can be accounted for under a simple population coding model and decision criterion, observing either of these behaviours experimentally does not necessarily discriminate between mechanisms (at least for orientation discrimination; Harrison & Bex, 2015). It may be fruitful to consider what such a letter-agnostic population coding model might predict for our experiments.

### Relevance to other investigations of distortions

How do our results fit with previous investigations of human perception of these two distortion types? We first consider BPN distortions. Our Experiment 1 revealed that distortion sensitivity is tuned to mid-range distortion frequencies (approximately 6–9 c/deg). Bex (2010) also found bandpass tuning for detecting BPN distortions introduced into one quadrant of natural scenes. Observers were maximally sensitive to distortions of approximately 5 c/deg, and these peaks were relatively stable for distortions centred at retinal eccentricities of 1.5, 2.8 and 5.6 deg. These estimates are at the lower bound of those we observe here. This might suggest that distortion detection sensitivity in letter stimuli peaks at higher spatial scales than detecting distortions of natural scene content. However, the results of Wiecek et al. (2014) imply that the peaks we observe will also depend on letter size, so it may not be generally meaningful to compare the peaks we observe to those of Bex (2010).

In Wiecek et al. (2014), letters of different sizes were presented foveally, and participants identified the letter after BPN distortion. Letter identification performance showed different tuning for distortion frequency at different letter sizes. Filtering with a peak frequency of 8 c/deg produced poorest identification performance for letters subtending 0.33 deg. These results fit with our data, if we assume that when a distortion is maximally detectable (peak sensitivities in our experiment) it maximally reduces letter identification (Wiecek et al. (2014)); the difference in letter size likely reflects a size scaling constant in detectability as letters move away from the fovea (Chung et al., 2002; Song, Levi, & Pelli, 2014).

What causes the bandpass tuning for BPN distortions? Potentially, sensitivity to whatever primitive feature dimensions are used to detect the distortions (e.g. contrast, curvature changes) also follow a bandpass shape. Note however that an analysis of the spatial frequency and orientation energy changes induced by distortions (Supplemental Material) reveals no obvious relationship to performance for those dimensions. Additionally, BPN distortions of sufficient amplitude (when the pixel shift exceeds half the distortion wavelength) will cause reversals in pixel positions, producing “speckling” at high frequencies but leaving the mean position of low frequency components unchanged (see for example Figure 1D, the highest amplitude distortions for the two highest frequencies). The bandpass tuning might reflect sensitivity to this speckling: detecting high frequency distortions requires detecting high frequency speckles (see also spectral analysis in the Supplemental Material), which are difficult to see in the periphery due to acuity loss^7^. Thresholds therefore rise again compared to mid-frequency distortions, which observers can detect well before speckling occurs. Experiment 1 also showed that when flankers are present, peak sensitivity shifts to higher frequencies than when flankers are absent. This could be because flanking letters selectively reduce sensitivity to position changes at lower spatial scales, or because flanking letters increase sensitivity to higher-frequency speckles. Given that there is no plausible mechanism that might support the latter possibility, we favour the former.

As to RF distortions, Wilkinson et al. (1998) measured thresholds for detecting RF distortions applied to spatially-bandpass circular shapes as a function of radial distortion frequency. They found that threshold amplitudes decreased as radial frequency increased as we do, but with a different pattern in which thresholds appeared to asymptote for higher frequencies. For RF1 patterns (which we do not test in our study), thresholds were ≈ 0.2, for RF2 patterns thresholds dropped to ≈ 0.01, and for higher frequencies (3–24 c/2*π*) thresholds asymptoted at an average amplitude of 0.003 (in the “hyperacuity” range). Thresholds in our data (Experiment 1 unflanked condition) were much higher (for example, average thresholds for our RF2 patterns were ≈ 0.15, which is about fifteen times higher than in their data). This is likely because distortions in our experiment were applied to more complex shapes (letters as opposed to bandpass circles) that were presented peripherally (whereas in Wilkinson et al’s experiment stimuli were nearer to the fovea). Nevertheless, there is little evidence that the asymptotic sensitivities in their results also hold in ours. This may be because the asymptote occurs for higher radial frequencies in the periphery, which conceivably reflects an interaction between the image content of our letter stimuli and the sensitivity of the peripheral retina. Dickinson et al. (2010, see also Dickinson, Mighall, Almeida, Bell, and Badcock (2012)) applied RF distortions to complex broadband images (faces) but did not characterise the radial frequency sensitivity function of these manipulations, so their results are not informative for this question.

### Caveats

The experiments in the present paper should be considered with a number of caveats. First, we measure performance for a single target-flanker spacing distance. While this distance was selected to be well within “Bouma’s law” for crowding, and we indeed find an influence of flanking letters, our data provide only a snapshot of the spatial interference profile for these stimuli. Successful models could also be expected to account for the spatial extent of crowding for letter distortions, and so measuring the spatial interference zones would be a useful experimental contribution. In the interests of brevity we leave those investigations to future studies.

Second, our results do not allow a direct comparison between the two distortion techniques. The frequency and amplitude parameters for each distortion type represent different physical image changes. Radial frequency distortions are highly correlated both tangentially and radially, whereas BPN distortions are not, and these correlations will interact with the original structure of the letter. Each distortion type produces different patterns of human sensitivity as a function of its distortion parameters. Therefore, the distortions and psychophysical results we present here define distinct physical shape changes that produce different patterns of sensitivity, providing a challenge for future accounts of shape perception.

Finally, the generality of our results should be considered with a degree of caution. The detectability of a given distortion will depend on the image content to which it is applied (for example, distorting a blank image region results in no image change). In our experiments we used only four target letter stimuli. This choice was motivated by the fact that our intention was not to quantify the visibility of distortions across a broad range of stimuli, but to investigate sensitivity in highly familiar simple patterns. Nevertheless, the research discussed above (Bex, 2010; Wiecek et al., 2014) corroborates the pattern of bandpass tuning we observe for the BPN distortions in our small set of letter stimuli, suggesting that this pattern applies more generally than just our limited stimulus set. As to RF distortions, we cannot say with any degree of certainty how the patterns of RF sensitivity we observe will generalise to new stimuli, because the previous investigations we are aware of either have not characterised distortion sensitivity as a function of frequency, or have done so in much simpler stimuli (see above).

### Other implications

The results of Wiecek et al. (2014) imply that the visibility and functional impairment caused by distortions originating in the retina (such as in metamorphopsia) will depend on viewing distance. Alongside the functional impact of these distortions for the patients in the real world, this result has important consequences for visual acuity testing in the clinic. Interestingly, patients with metamorphopsia often fail to notice their distortions in the real world (Wiecek, Lashkari, Dakin, & Bex, 2015) and even when tested with artificially-regular stimuli (Crossland & Rubin, 2007; Schuchard, 1993; Wiecek et al., 2015). “Filling-in” processes (Crossland & Rubin, 2007) and binocular masking (Wiecek et al., 2015) undoubtedly contribute to this insensitivity. To the extent that the results we report here are generalisable (see above), they (along with Bex, 2010) offer an additional explanation for why patients with metamorphopsia often fail to notice their distortions: in the real world, distortions caused by retinal disease will often be crowded by cluttered visual environments.

## Conclusion

Taken together, the pattern of results presented here provide a challenge for models of 2D form processing in humans. A successful model of form discrimination would need to explain sensitivity to two distinct distortion types, the dependence of distortion sensitivity on flanking letters, and the dependence on the type of flanking letters (distorted flankers reduce sensitivity). Directly comparing the BPN and RF distortions would require an image-based similarity metric that captured the perceptual size of the distortions on a common scale. One test of such a similarity metric would be to rescale the results of the BPN and RF data reported here such that the different sensitivity patterns as a function of distortion frequency overlap (assuming that they are detected by a common mechanism). We have provided our raw data and images of the stimuli used in these experiments (http://dx.doi.org/10.5281/zenodo.159360) to facilitate future efforts along these lines.

## Acknowledgements

Designed the experiments: TSAW, ST, FAW, MB. Programmed the experiments: ST, TSAW. Collected the data: ST, TSAW. Analysed the data: TSAW, ST. Wrote the paper: TSAW. Revised the paper: ST, FAW, MB. We thank Peter Bex and William Harrison for helpful comments on the manuscript.

## Supplemental material

### Difficulty of individual target letters

Because the effect of an image-based distortion depends on the image content, we note that performance varied slightly according to the target letter (Figure 6). On average across observers, it was easier to detect distortions applied to the letters K and H than the letters D and N, for both distortion types. Note however that the comparisons in Figure 6 conflate distortion sensitivity and response bias. Because each letter is presented on every trial (with the distortion applied to only one of the letters), an observer with a bias to choose a particular letter when in doubt (irrespective of its location) would also serve to raise proportion correct performance (or thresholds). Thus, biases that are consistent across observers could also produce differences in letter performance. Measuring sensitivity to distortions in each letter while eliminating bias would require a forced-choice on individual letters (e.g. which of these “K”s is distorted?). Nevertheless, we find this possibility unlikely because it would require observers to identify the location of their preferred letter and respond accordingly—it therefore seems more plausible that response biases would occur for response locations rather than for letter identities. Another possible explanation for different letter sensitivities is revealed by considering that the advantage for K and H appears larger in flanked than unflanked conditions. These effects could depend on the relative similarity of the target letters and the four flanking letters, which have been shown to influence letter identification under crowded conditions (Bernard & Chung, 2011; Hanus & Vul, 2013).

**Figure 6.**
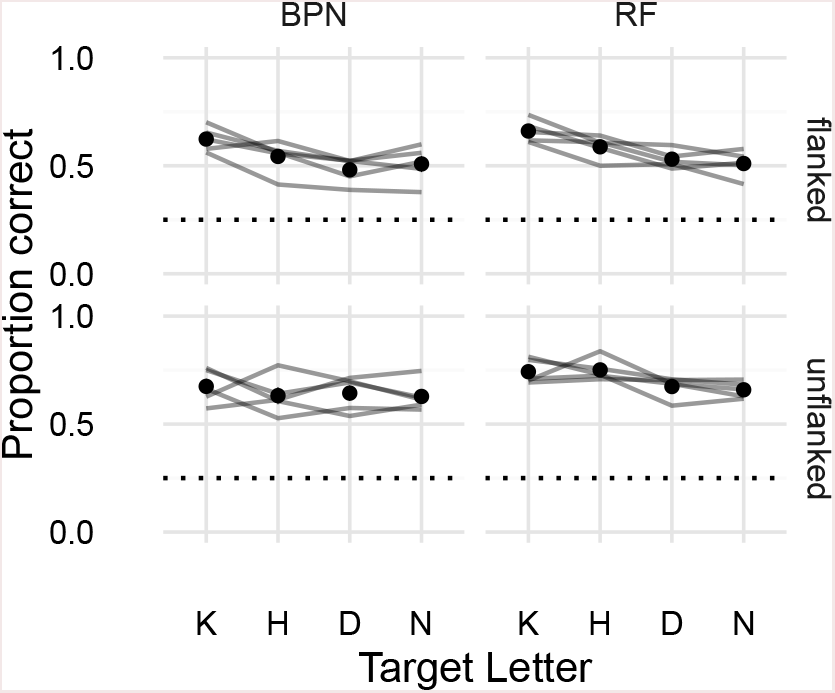
Performance for each target letter in each distortion type and flanking condition. Points show the average proportion correct across observers (error bars show ±1 SE) for each target letter in each distortion type, averaged over frequencies and amplitudes. Lines link the performance of individual observers. The letters K and H show slightly higher performance than D and N, for both distortion types, and this trend appears slightly stronger for flanked than unflanked trials. This could reflect an interaction between letter shape and distortion (i.e. it is easier to discriminate distortions applied to the letter K), differential similarities of the target and flanking letters, or biases in preferred letter irrespective of location.

### Analysis of spatial frequency and orientation spectra

To gain insight, into the physical changes caused by the letter distortions that may underlie the results we observe, we examined how the different distortions change the spatial frequency and orientation energy spectra of the stimuli. If observers were able to perform the task by simply detecting spectral changes in the target letters, then we would expect the physical changes caused by the distortions to mirror the patterns of sensitivity from Experiment 1. Specifically, for BPN distortions we should observe a bandpass tuning of the relevant dimension (peaking for middle distortion frequencies) whereas for RF distortions we should observe a spectral change that increases with distortion frequency.

We computed the Fourier amplitude spectrum of each target letter image (92×92 pixels), then calculated the radial energy (averaging over angle) and angular energy (averaging over radius) by applying Gaussian sliding windows (using the `spectral_analysis` function from Psyutils v1.3.1: http://dx.doi.org/10.5281/zenodo.159360). These correspond to the spatial frequency and orientation energies respectively. We performed this operation for the undistorted target letters, and for letters distorted with BPN and RF distortions with frequencies as in Experiment 1 and for three distortion amplitudes: the amplitude corresponding to the average threshold for the unflanked condition, the flanked condition, and for the maximum distortion we applied in Experiment 1. Within each combination of conditions we generated 15 unique distorted letters (i.e. with different noise patterns for BPN and different phases for RF) in order to capture the average effect of the distortions.

For spatial frequency energy, high-frequency BPN distortions increased the contrast energy in high spatial frequencies (≈ 8–16 c/deg; see Figure 7) compared to the undistorted letter, for all letters. Even if we assume that these frequencies are easily detectable at the ≈ 8 degrees of retinal eccentricity used in our study, it seems unlikely that observers benefit from this increased contrast energy because thresholds for these conditions were higher than those for lower frequency distortions (i.e., BPN distortion sensitivity follows a bandpass shape). High frequency RF distortions also increased high spatial frequency energy (Figure 8), but not as much as for BPN distortions (Figure 9). RF distortions of 5 and 8 c/2*π* also increased frequencies in the mid SF range (4–8 c/deg; more easily seen in Figure 9), but again this increased contrast energy appears unrelated to psychophysical performance. Considering these results across the two distortion types, it seems unlikely that changes in spatial frequency energy could underlie human performance in our experiment. For example, if observers were using the increase in high spatial frequency energy to perform the task, then we would expect thresholds in the high frequency BPN conditions to continue declining; instead they increase again.

**Figure 7.**
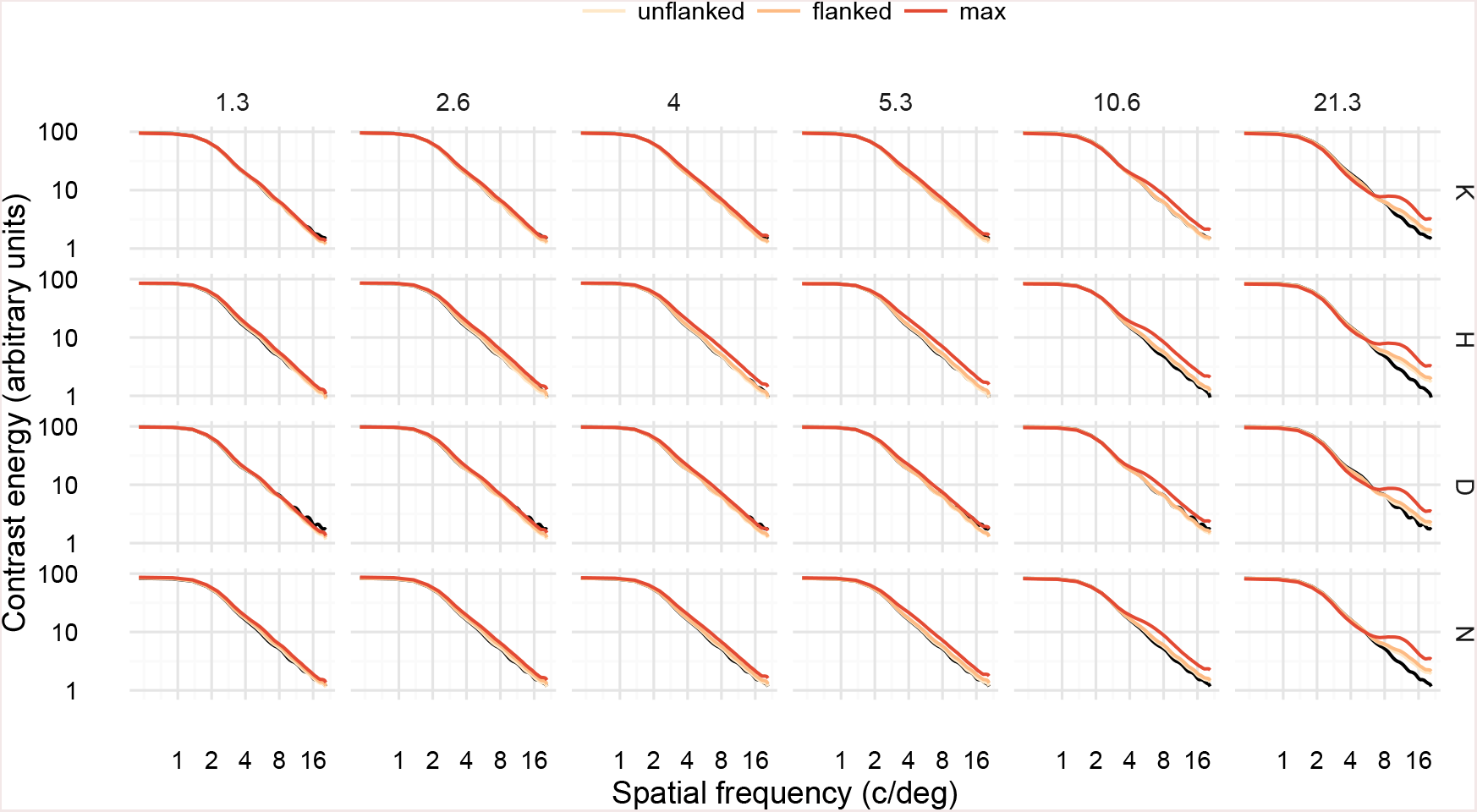
Spatial frequency energy in letter stimuli with BPN distortions. Panels are arranged by target letter (rows) and distortion frequency (columns). Black curves show the spectrum of the undistorted letter; coloured curves show the spectra for letters distorted at threshold levels for unflanked and flanked conditions, as well as the maximum distortion used in Experiment 1.

**Figure 8.**
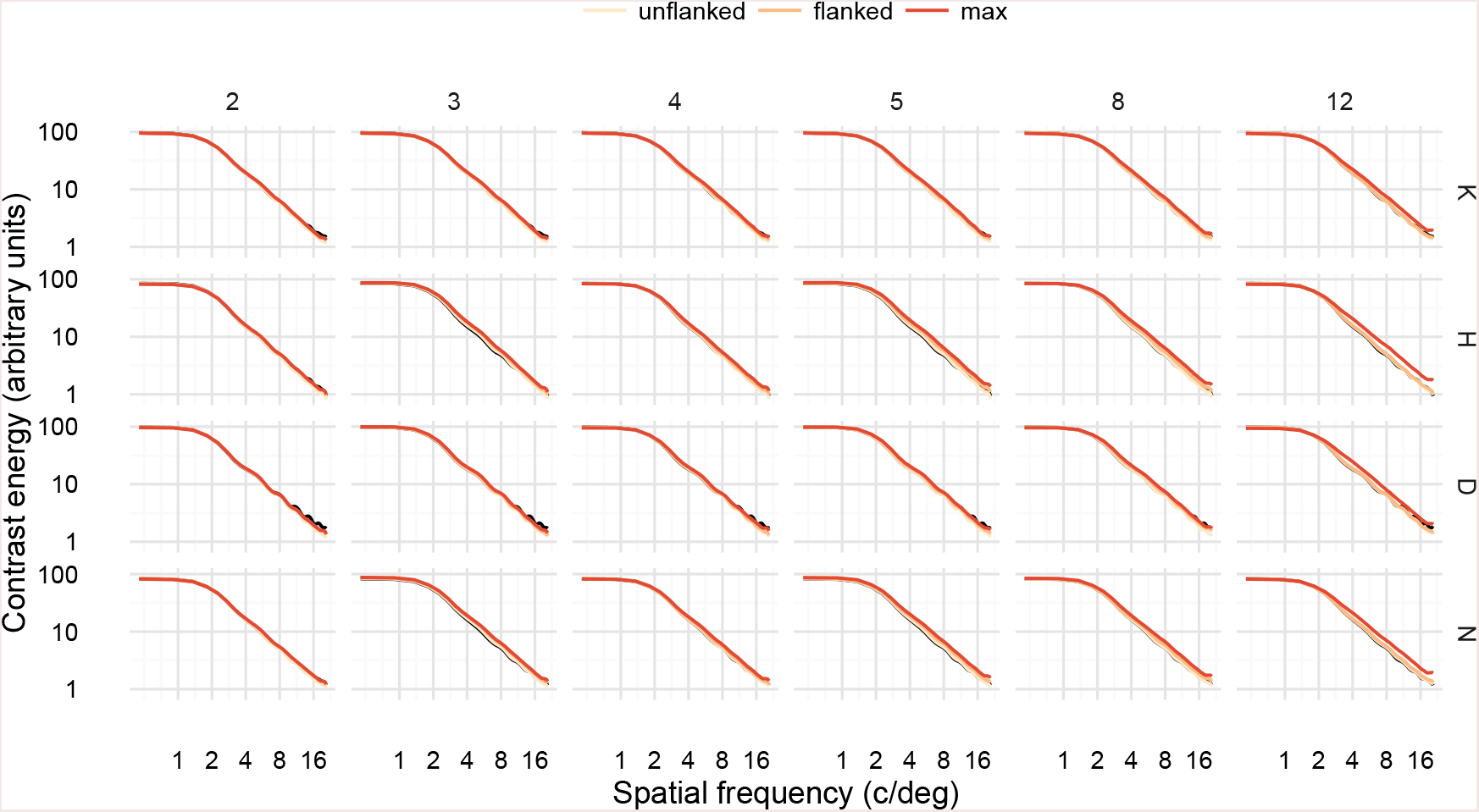
Spatial frequency energy in letter stimuli with RF distortions. Plot elements as in Figure 7.

**Figure 9.**
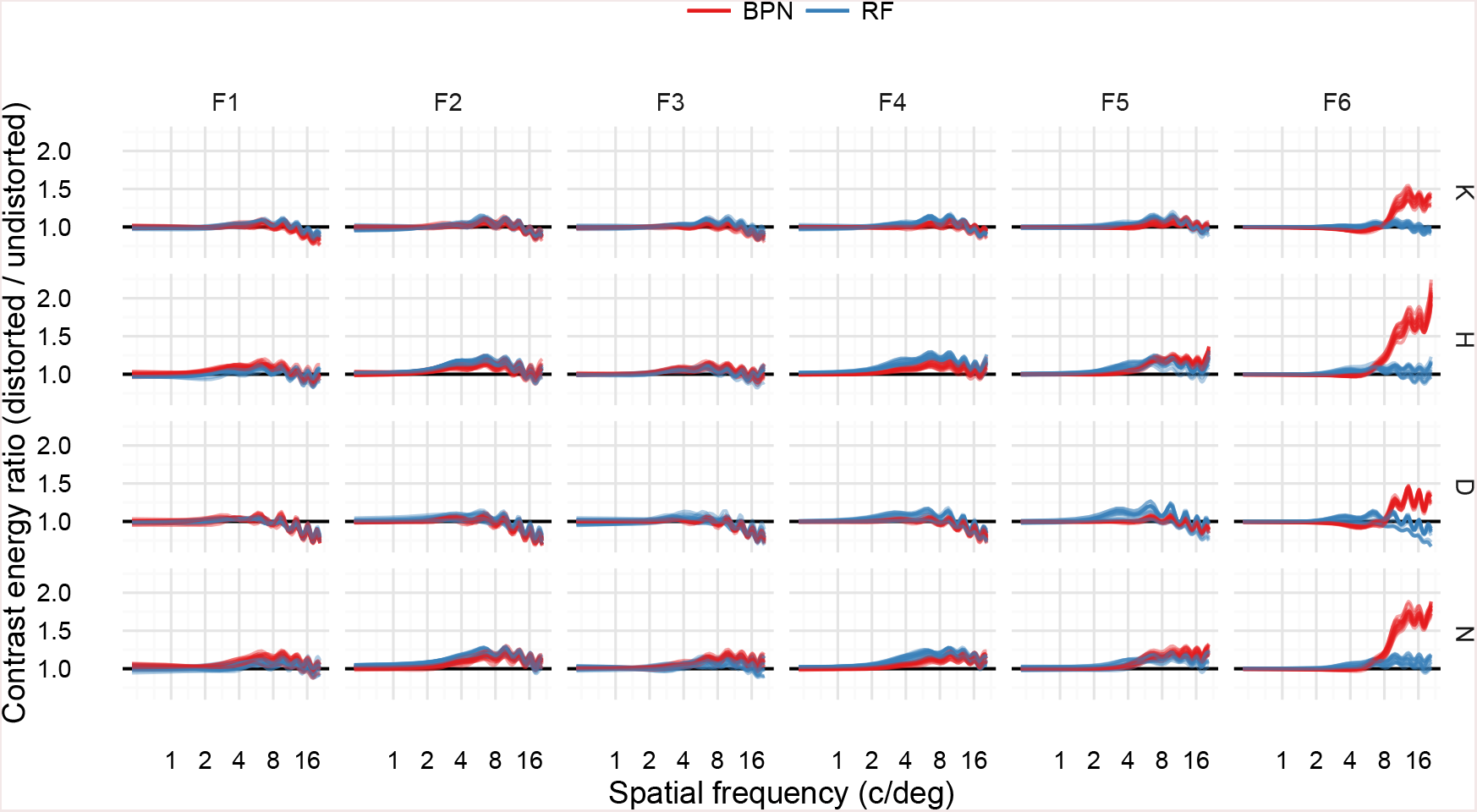
Ratio of spatial frequency energy between undistorted and distorted letters, for BPN and RF distortions, at amplitudes corresponding to average flanked thresholds. Each faint line shows a unique distortion. Distortion frequencies have been categorised into the lowest (F1) to highest (F6) shown for each distortion type. High-frequency BPN distortions increase contrast energy at high spatial frequencies.

For orientation energy, both BPN (Figure 10) and RF (Figure 11) distortions have the effect of increasing energy at all orientations relative to the original letter stimulus (making the distribution of orientation energy less peaked) as distortion frequency increased. This effect is much more pronounced for the BPN distortions compared to the RF distortions (Figure 12) at the highest distortion frequency. Again, this pattern of physical stimulus changes holds no obvious relationship to psychophysical performance. If observers used changes in the orientation energy to detect the distorted letter, then for BPN distortions we would expect these changes to be greatest for F4 or F5 (5.3 or 10.6 c/deg) for the flanked condition, and for RF distortions we would expect the energy change to increase with distortion frequency. Instead, the highest BPN frequency has the largest relative effect on orientation energy, and if anything the effect of RF distortions are greatest for middle distortion frequencies (F4 or F5; Figure 12).

**Figure 10.**
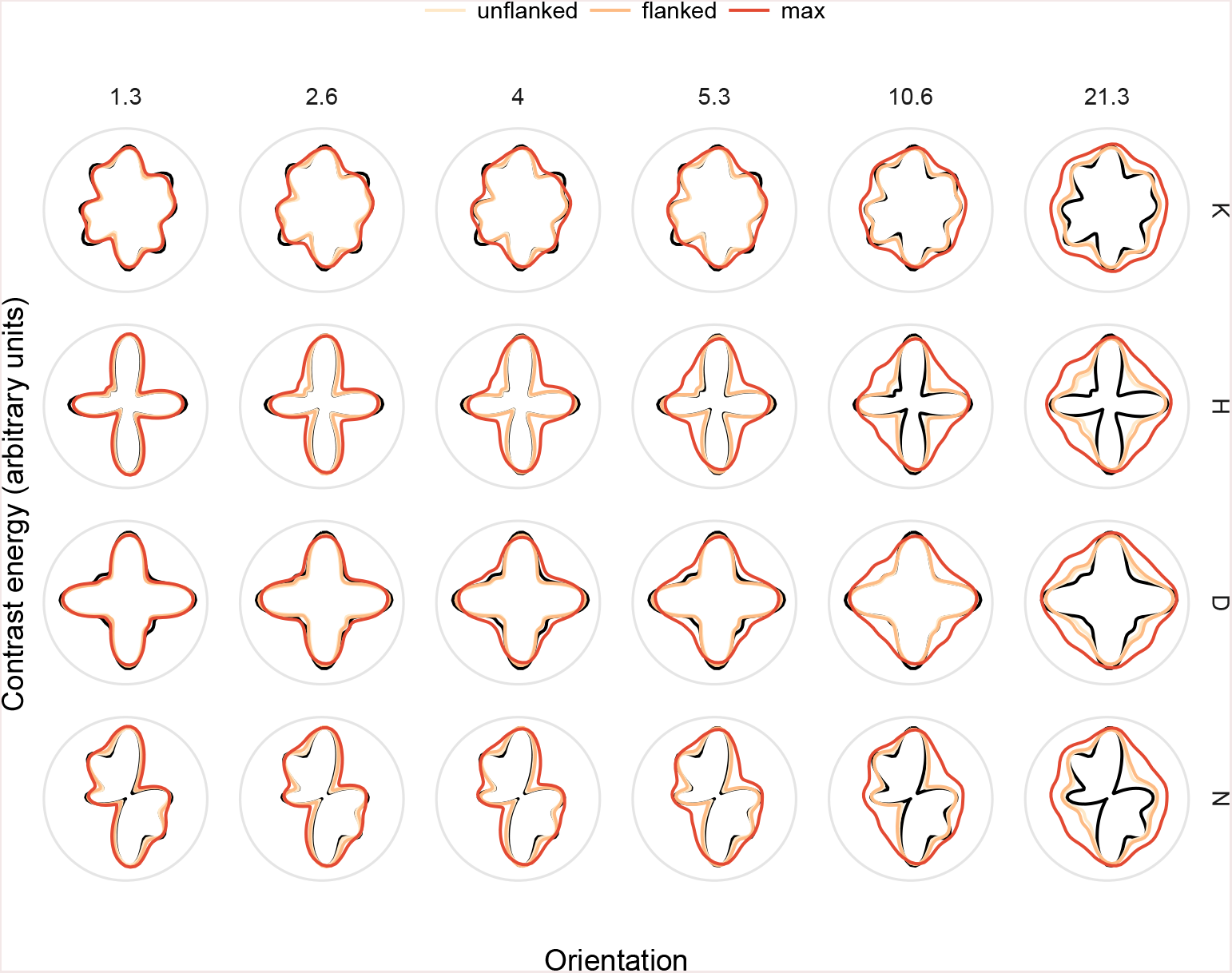
Orientation energy in letter stimuli with BPN distortions. Polar coordinates have been rotated relative to the orientation of the raw amplitude spectra so as to be more intuitive. To gain an intuition for the orientation, consider the original spectrum for the letter “N”. Imagining an upright “N”, one can see most energy at vertical orientation and also on the diagonal corresponding to the diagonal stroke in the letter. Panels and plot elements as in Figure 7.

**Figure 11.**
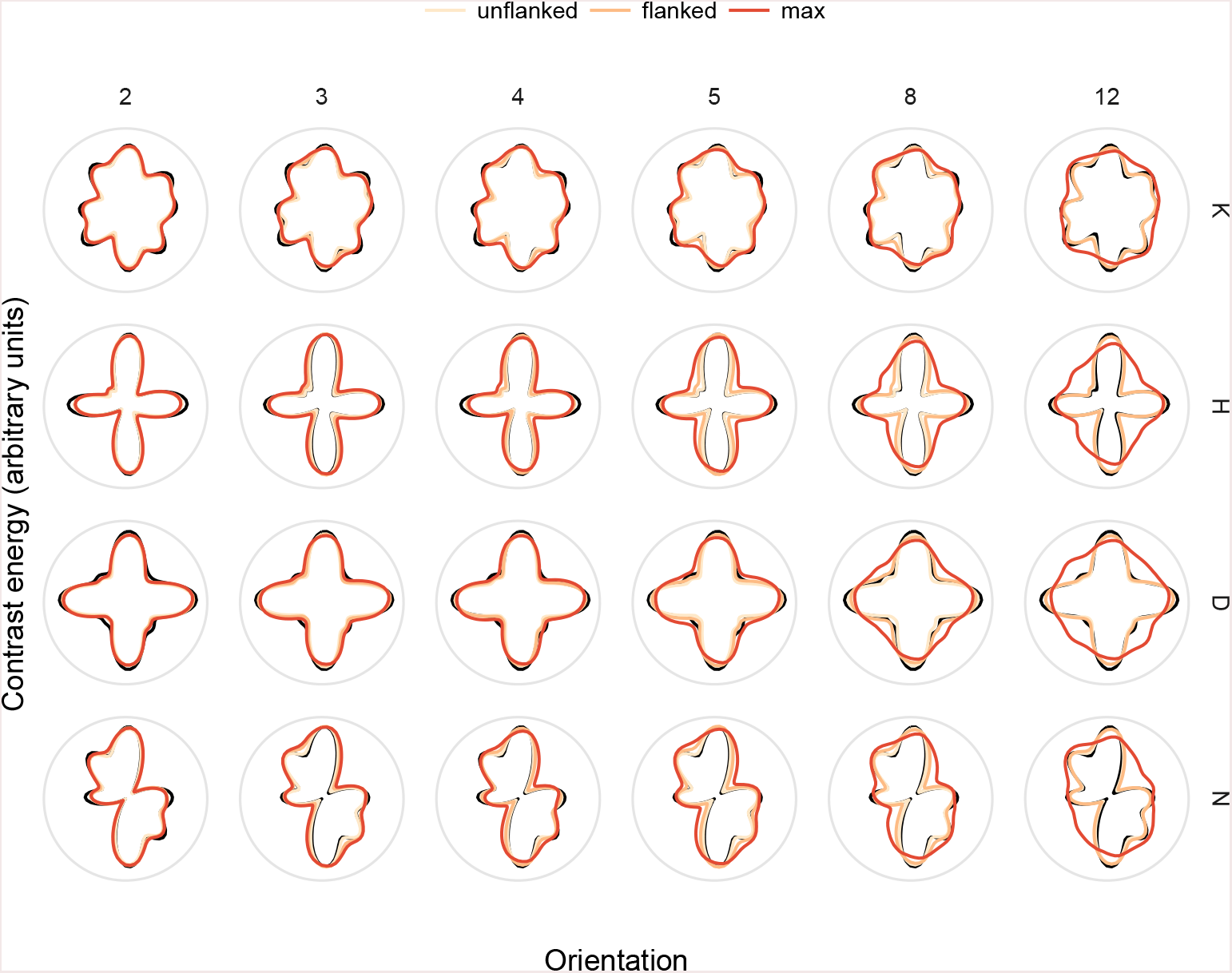
Orientation energy in letter stimuli with RF distortions. As for Figure 10.

**Figure 12.**
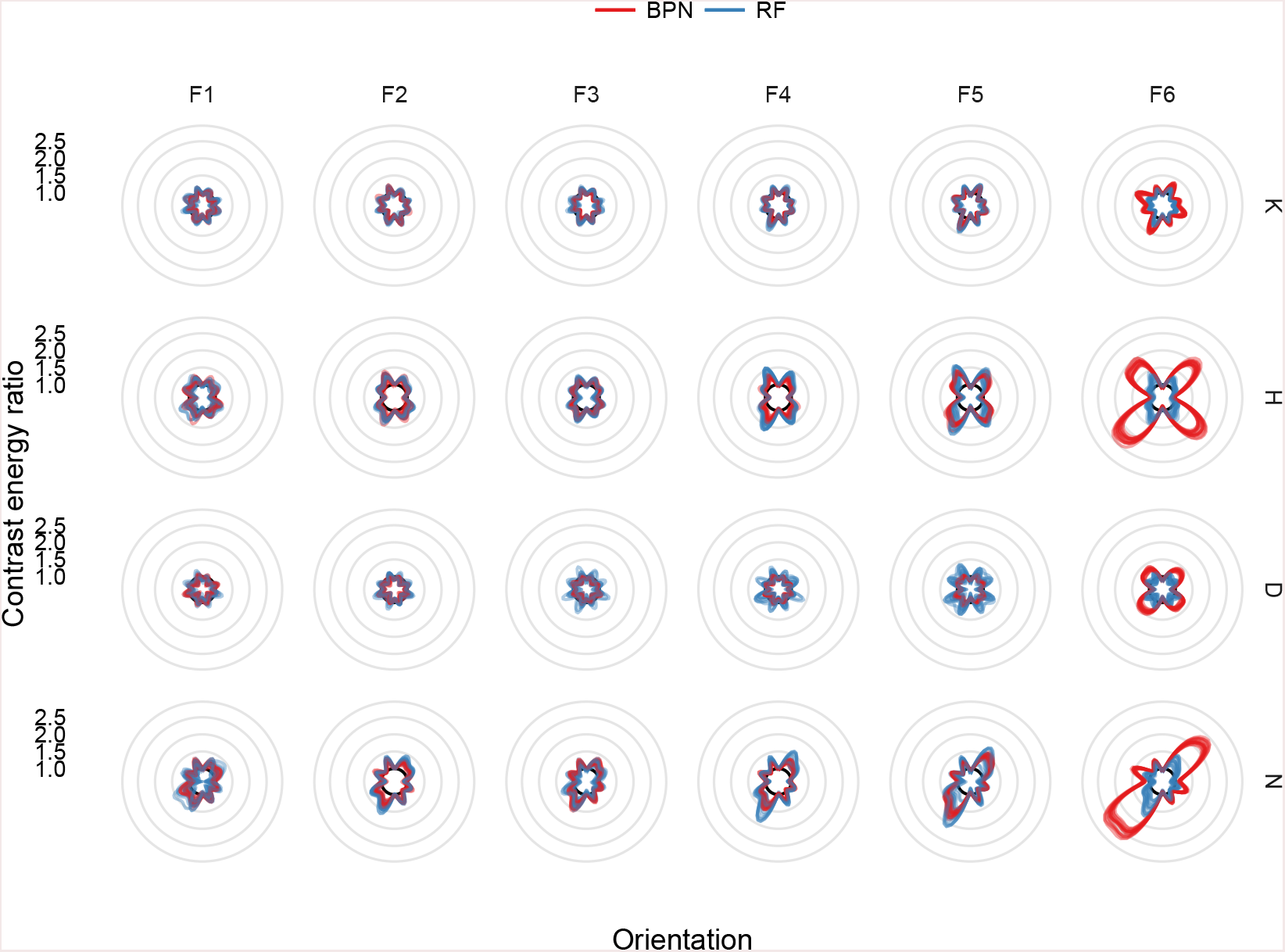
Ratio of orientation energy between undistorted and distorted letters, for BPN and RF distortions, at amplitudes corresponding to average flanked thresholds.

Finally, we can consider whether differences in frequency or orientation energy might underlie the slightly different performance for target letters, in which observers were more sensitive to K and H than D and N for both distortion types (Figure 6). The largest changes in both spatial frequency and orientation energy are observed for H and N (Figures 9 and 12), which provides little evidence one way or the other. At least, there is no strong evidence that changes in spatial frequency or orientation energy drive differential performance for these target letters.

### Clutter metric analysis

In an attempt to provide some quantitative basis for our speculations about display complexity as an account for the results of our second experiment, we here apply two metrics for “clutter” to our stimulus displays (Rosenholtz et al., 2007). The first metric, *feature congestion,* is a multiscale measure of the covariance of three features: the luminance contrast, orientation and colour in a given input image (since our images are greyscale, the contribution of colour will be minimal). The second metric, *subband entropy,* is determined by the bitdepth required for wavelet image encoding, expressed as Shannon entropy in bits. For both metrics, higher values are associated with more “cluttered” images. These metrics have been shown to be predictive of aspects of visual search performance across a variety of domains (Asher et al., 2013; Henderson et al., 2009; Rosenholtz et al., 2007).

We used the publically-available code from Rosenholtz et al. (2007, see https://dspace.mit.edu/handle/1721.1/37593). We analysed all the images used in both experiments. In all cases we report the scalar clutter metric (averaged over space in the image to give one number per stimulus display). Figure 13 shows the results for Experiment **1.** While both metrics qualitatively reproduce the effect of adding flanking letters (clutter increases in the “flanked” condition), neither metric produces the qualitative pattern of results as a function of frequency (bandpass patterns for BPN distortions or decreasing clutter as frequency increases for RF distortions), at least at the scale of the increase caused by adding flanking letters.

**Figure 13.**
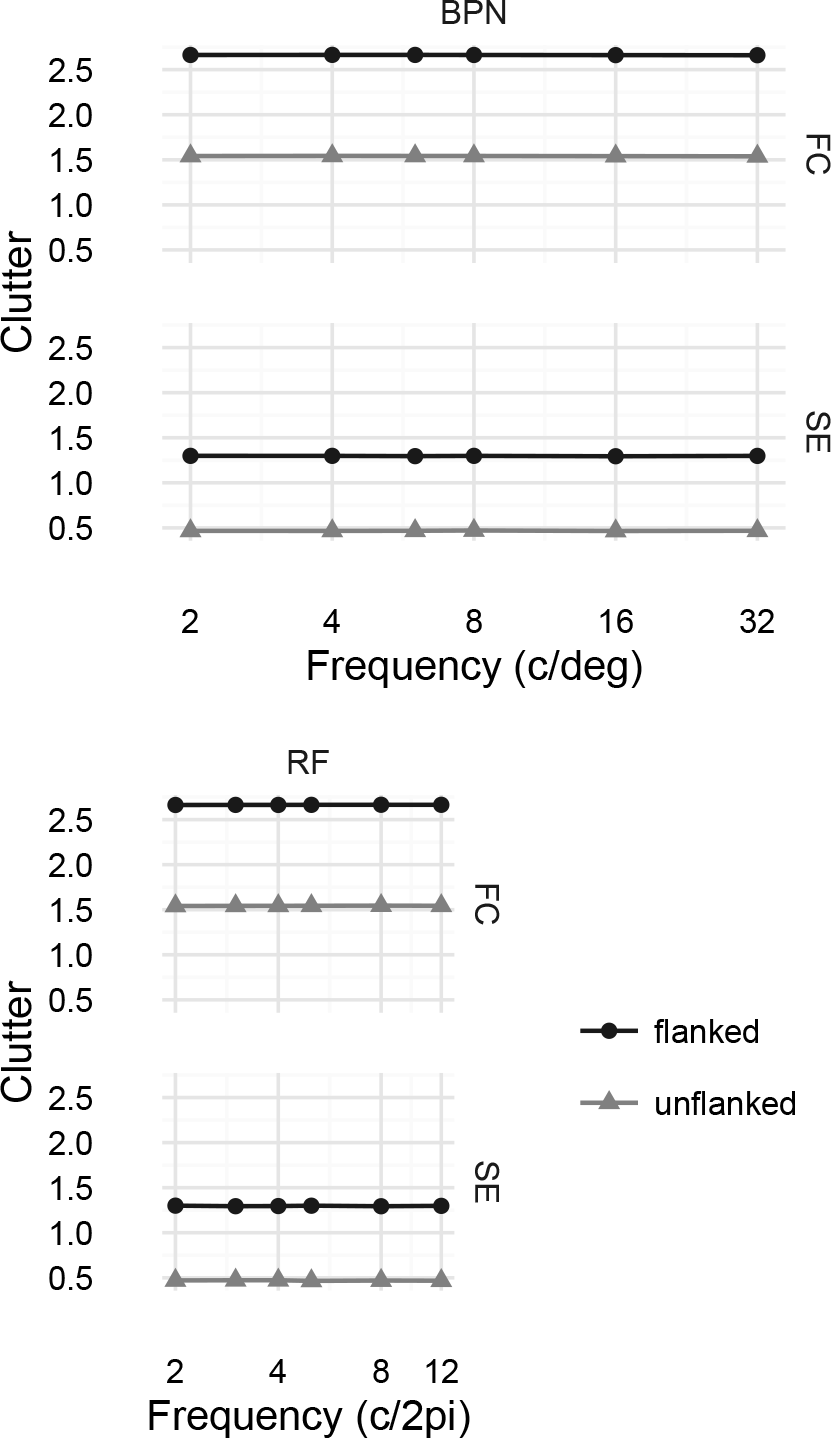
Clutter analysis of Experiment 1. Feature congestion (FC) and subband entropy (SE) clutter metrics for our BPN and RF distortion images as a function of distortion frequency and flanked / unflanked display. Points linked by lines show the value of the metric averaged over distortion amplitudes (which had little effect on clutter at this scale), letters and unique stimuli. Flanking letters substantially increase visual clutter.

In Figure 14 we plot only the flanked condition for the highest distortion amplitude. While one could optimistically see interesting patterns in the means of the FC clutter metric at this scale (apart from the dip at 4*c*/2*π*, RF clutter increases with frequency, whereas BPN clutter shows hints of a bandpass shape, albeit at a lower peak than the human data), the large variance for individual images provides at best weak evidence for any correspondence with the human data from Experiment 1.

**Figure 14.**
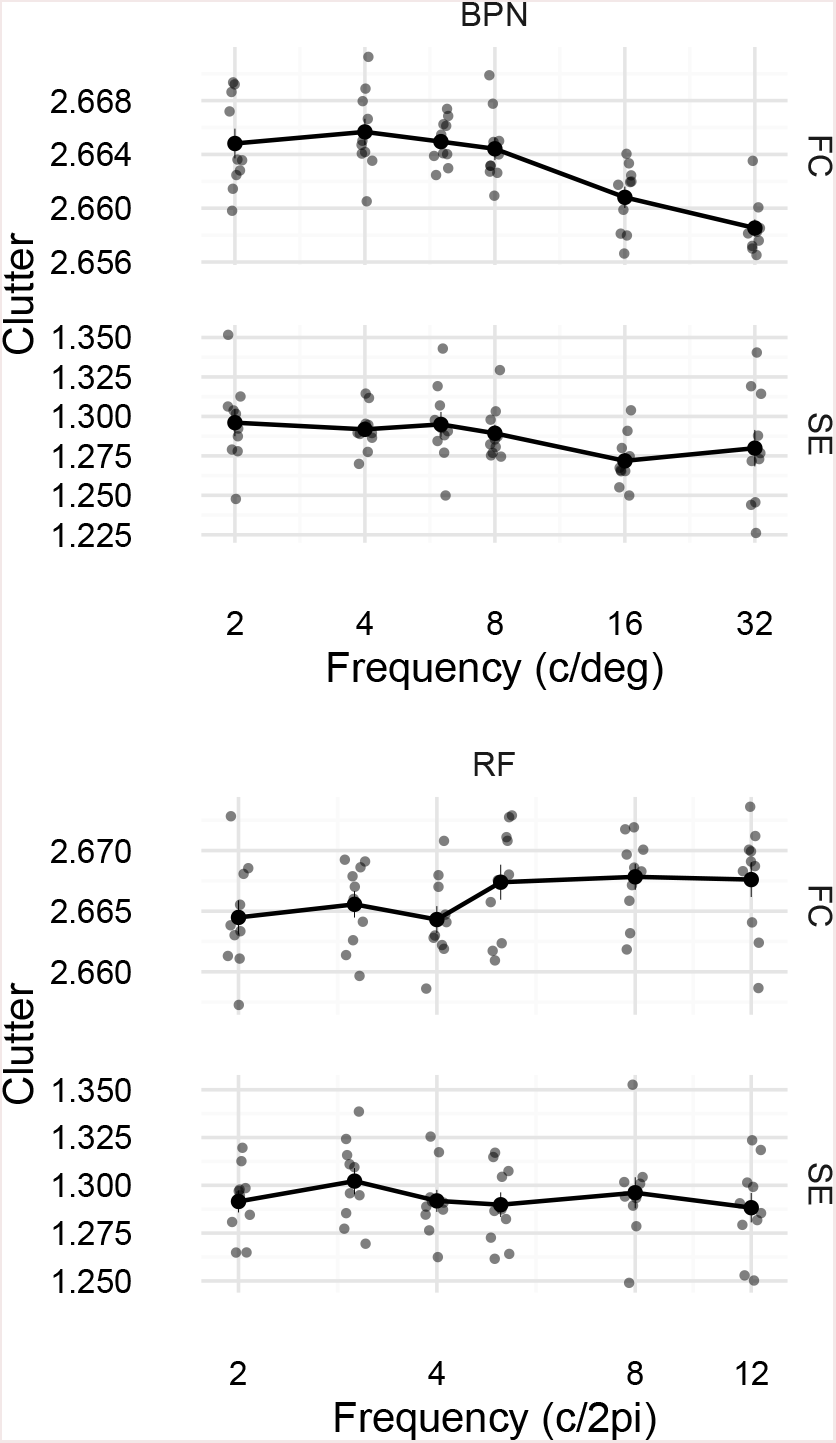
Clutter analysis for Experiment 1, flanked condition at the highest distortion amplitude. Each small semitransparent circular point represents one image from the experiment (these have been jittered on the x-axis to reduce overplotting). Filled circles linked by lines represent mean clutter (±1 SEM).

Can the clutter metrics account for the influence of the number of distorted flankers (Experiment 2)? The change in clutter caused by applying distortions to zero, two or four flankers is shown in Figure 15 (stimuli from Experiment 2a). The means of the feature congestion metric show a similar pattern of results as humans. This could imply that when detecting a distorted target amongst distorted flankers (Experiment 2a), more clutter (caused primarily by the four distorted flankers, which are uninformative about the target location) is associated with worse performance. While this could be taken as quantitative support for our suggestion that performance in Experiment 2 can be explained by display complexity, we would advise not to take this interpretation too seriously for the following reasons: first, the scale of the clutter changes here is tiny compared to the influence of adding flanking letters in the first place (whereas human threshold changes are both very robust). Second, as for the data for Experiment 1 (Figure 14), there is a large amount of variation in the individual images (presumably related to different configurations of target and flanking letters rather than effects of distortions per se).

**Figure 15.**
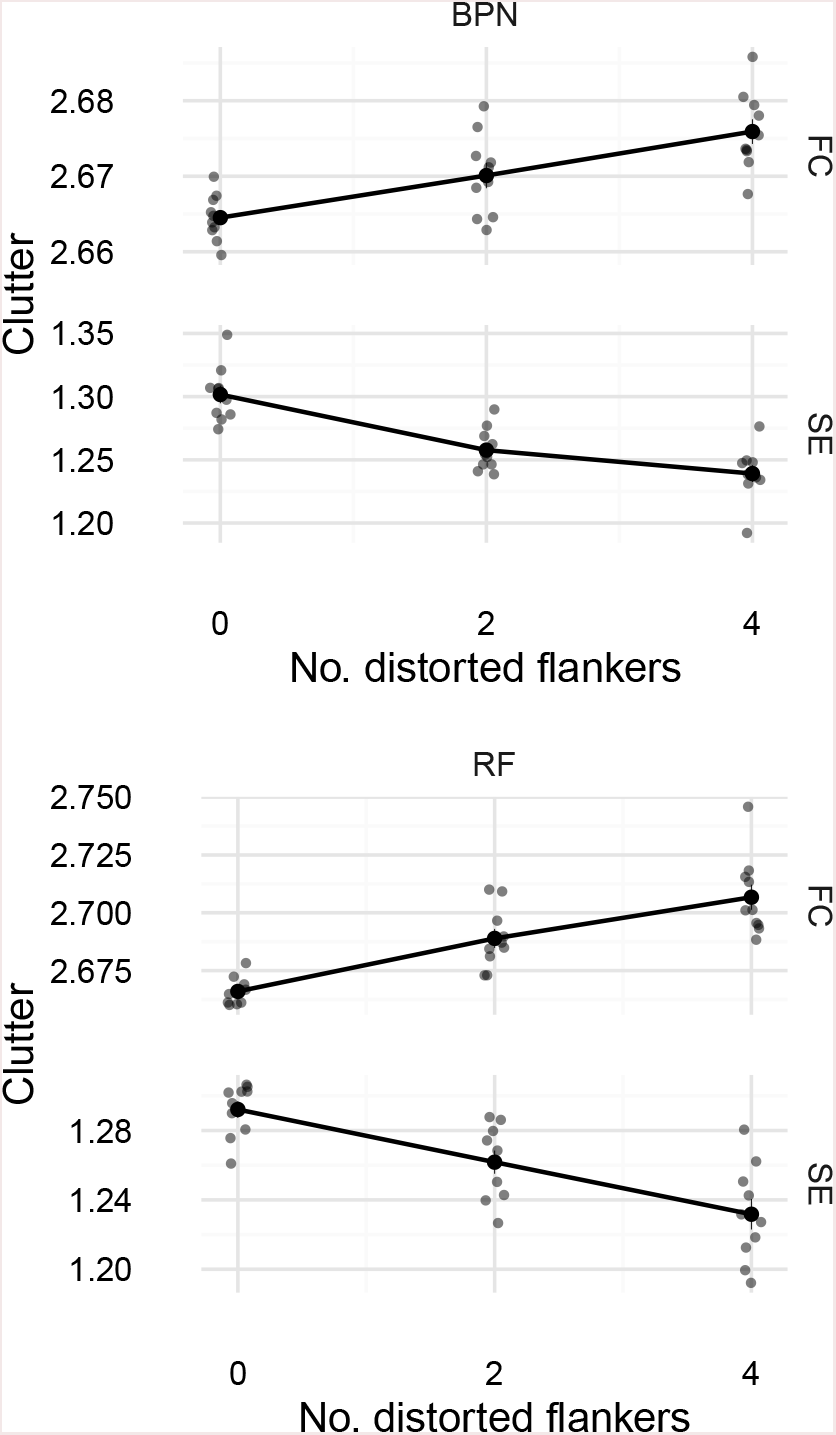
Clutter analysis of Experiment 2a. Results are shown for the highest distortion amplitudes only. Plot elements as in Figure 14.

Taken together, these results suggest that while image-based clutter metrics such as feature congestion or subband entropy account for the difference between the “unflanked” and “flanked” conditions of Experiment 1, one would need to find a more appropriate measure of complexity, or at least apply some transform to feature congestion, to capture the more subtle dependencies in our data.

### Examples of stimuli

Here we provide additional examples of distortions applied to different target letters. Figure 16 shows examples for BPN distortions applied to each letter, and Figure *17* show example letter distortions for the RF method.

**Figure 16.**
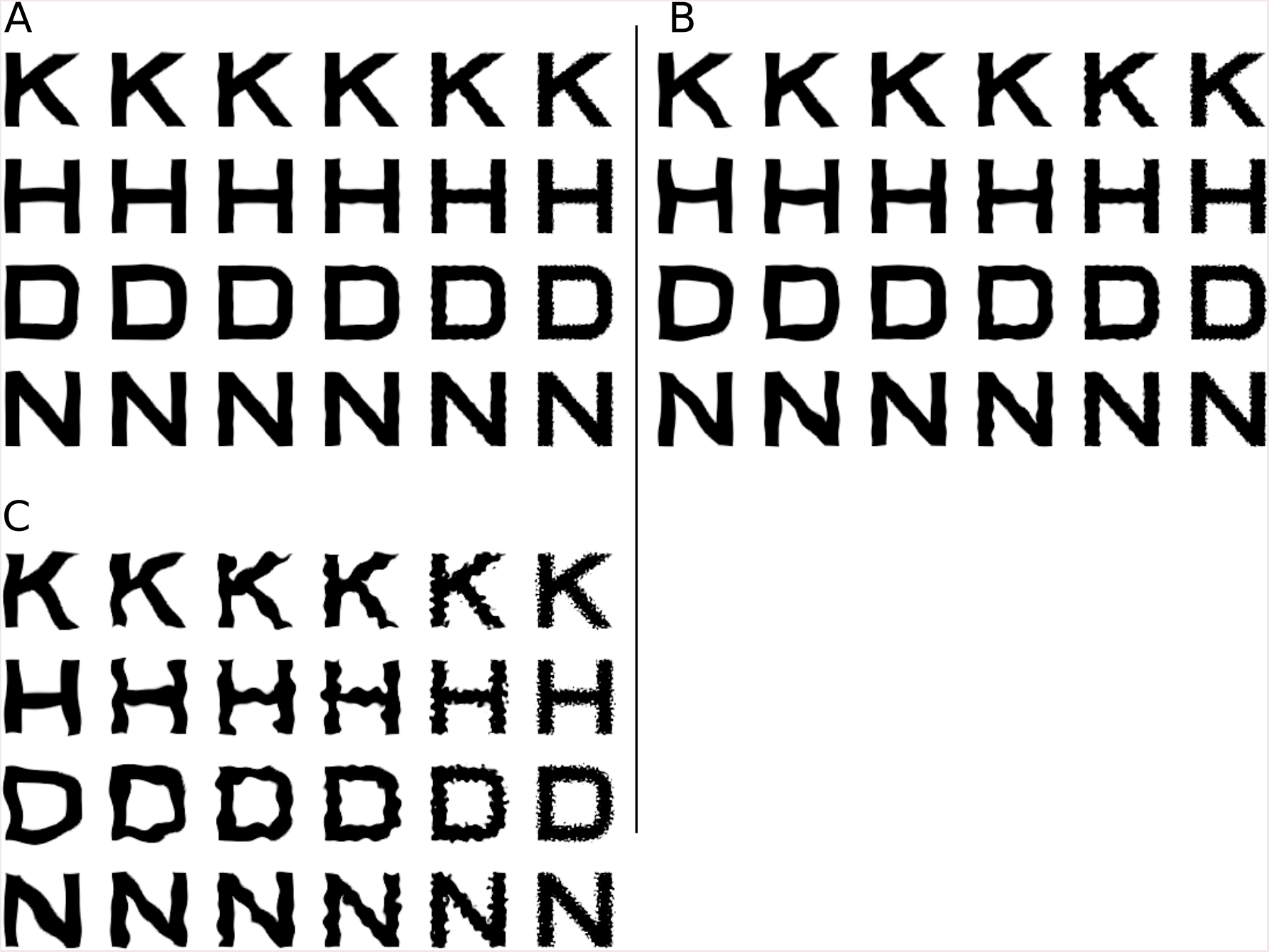
Examples of Bandpass Noise distortions. **A:** Letters (rows) distorted at the averaged unflanked threshold from Experiment 1. Columns show increasing distortion frequencies. **B:** Letters distorted at the averaged flanked threshold from Experiment 1. C: Letters distorted at the maximum distortion used in Experiment 1.

**Figure 17.**
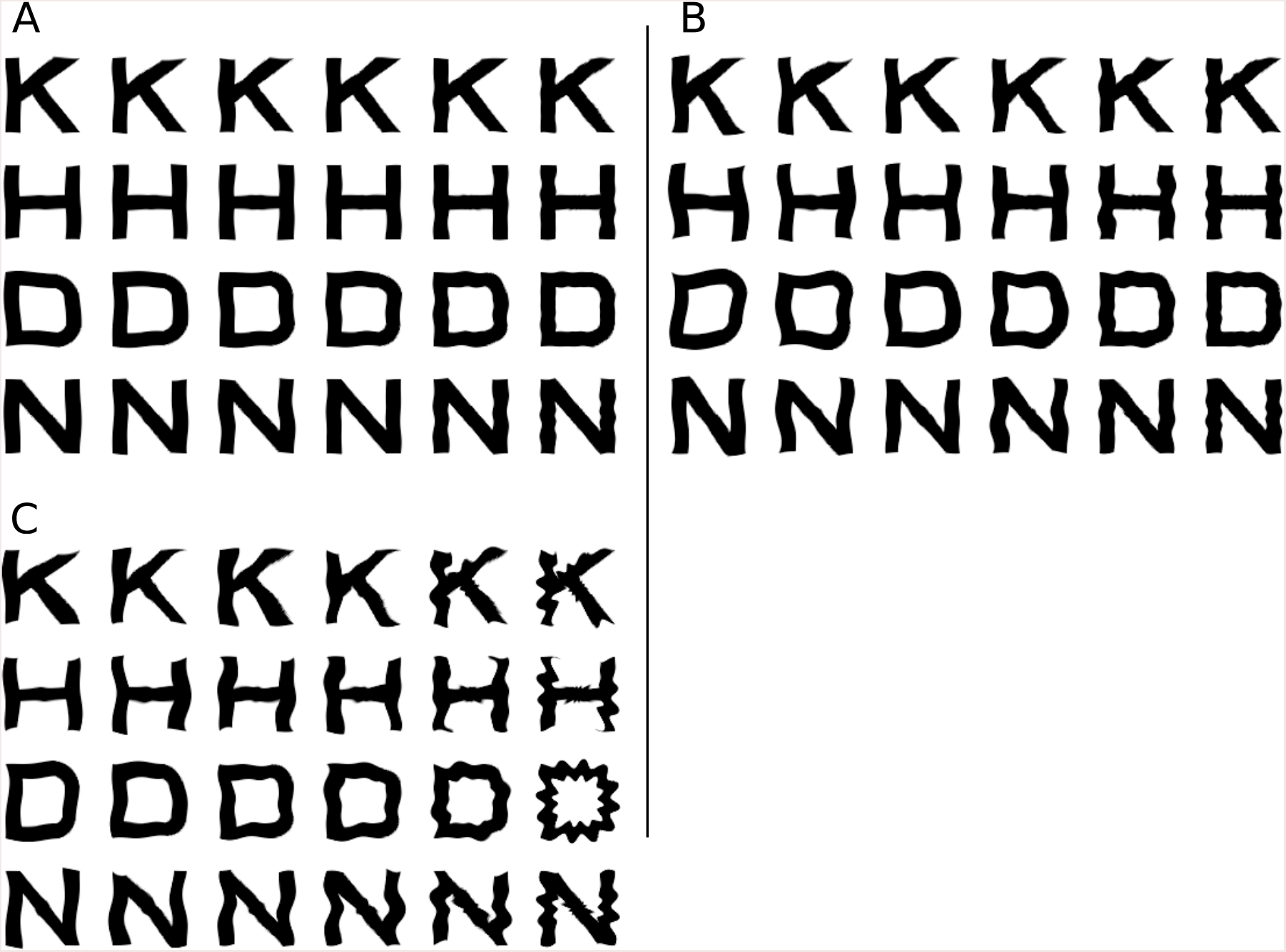
Examples of radial frequency distortions. Panels as in Figure 16.

## Footnotes

1 We generated stimuli for eight amplitudes but adjusted the sampling range after pilot testing to better sample the range of performance. All observers have done some trials at all amplitudes.

2 Any practice effect should therefore improve performance in the flanked condition (this is not what we found).

3 Note that these four-parameter functions are rather unconstrained by only six data points, and are intended as a rough guide to the patterns in the data rather than a definitive statement about tuning. A more robust estimate could be gained by fitting a mixed effects model.

4 While for the linear model we could directly compare intercepts and slopes, the area provides a simple measure that also accounts for different curvature in the Gaussian model.

5 For the BPN distortions, the average threshold in the flanked condition for Experiment 1 at a frequency of 2 c/deg was 0.06 (SD = 0.01), whereas with four distorted flankers (Experiment 2a.) the average threshold at the same frequency was 0.13 (SD = 0.03; a factor of 2.1 times larger). Similarly, in Experiment 1 the threshold for RF distortions at 4 c/2*π* was 0.19 (SD = 0.03) whereas the same frequency with four distorted flankers in Experiment 2a was 0.61 (SD = 0.01; a factor of 3.3 times larger).

6 We would like to credit a discussion with Daniel Coates that resulted in this (post-hoc) account of our data

7 We credit Peter Bex for pointing out the likely relevence of speckling to the observed tuning.

